# Shedding dynamics of the Ross seal (*Ommatophoca rossii*) whiskers investigated through carbon and nitrogen bulk stable isotope composition

**DOI:** 10.1101/2025.11.04.685852

**Authors:** Nicolas Séon, Trevor McIntyre, Maëlle Connan, Marthán N. Bester, Horst Bornemann, Sarah Bury, Mila B. Geldenhuys, Candice B. Lewis, Mia Wege

## Abstract

Understanding the trophic ecology of pinnipeds is essential to define their role in ecosystems and anticipate their responses as top predators to environmental changes. Incremental analyses of carbon (*δ*^13^C) and nitrogen (*δ*^15^N) stable isotopes along their whiskers provide valuable records of their foraging habitat and trophic level over time. The effectiveness of this approach relies on knowledge of species-specific whisker shedding dynamics and growth rates. We investigated the shedding dynamics of Ross seal (*Ommatophoca rossii*) whiskers and attempted to evaluate the time captured in these tissues by combining whiskers’ bulk *δ*^13^C and *δ*^15^N incremental measurements with satellite tracking data, collected after whisker growth. Our tracking data are consistent with previous studies, in that Ross seals migrate post-moult between the marginal ice zone and the Antarctic Polar Front. Stable isotope profiles of whiskers did not show the typical decrease in *δ*¹³C and increase in *δ*¹⁵N values at the whiskers’ tip that would be expected if shedding coincided with the annual fur moult, when fasting typically occurs. The occurrence of similar isotope variations, which were offset between the left and right whiskers of the same individual in relation to distance from the muzzle, further suggest asynchronous, non-seasonal whisker shedding. Our inability to identify known cyclic phenological events (i.e., fasting periods or seasonal migrations) prevented the determination of the average growth rate of whiskers. The additional data on whisker shedding dynamics of Ross seals is a valuable first step in support of future ecological studies based on the whiskers of this species.

## 1. INTRODUCTION

Time–depth recorders, accelerometers, stomach temperature sensors and animal–mounted cameras have been employed to gain insight into the foraging behaviour of marine mammals (Le Boeuf et al. 2000, Bradshaw et al. 2004, Molinet et al. 2025). However, the trophic ecology of some marine mammal species remains poorly understood, especially for species that inhabit the ice-covered polar oceans where it is difficult to access them and to set up these tools (Southwell et al. 2012). The Ross seal (*Ommatophoca rossii*), from the Phocidae family (Berta et al. 2018, i.e. true / earless seals), is one of these hard–to–reach marine species. Highly variable population estimates (between 20,000 and 254,500 individuals, Erickson & Hanson 1990, Bengtson et al. 2011, Curtis et al. 2011) and a paucity of data about their movement behaviour, reproduction, foraging ecology and diet makes it currently the only Specially Protected Species under the Antarctic Treaty System (SCAR 2007, Hughes et al. 2024). The lack of biological information about Ross seals is largely due to their inaccessibility as they spend most of their time at sea (Blix & Nordøy 2007, Arcalís-Planas et al. 2015, Wege et al. 2021, 2023). Therefore, opportunities for direct sampling are limited to brief periods when Ross seals haul out on the largely inaccessible Antarctic seasonal pack ice, primarily during late October – late November for pupping and nursing (Solyanik 1964, Thomas et al. 1980, Southwell et al. 2003, Bester et al. 2021) and in January – February in the densely consolidated pack ice for moulting (Blix & Nordøy 2007, Arcalís-Planas et al. 2015, Wege et al. 2023).

Bulk stable isotope analyses of animal tissues provide a complementary approach for gaining insights into Ross seal trophic ecology by determining their foraging habitats through their carbon stable isotope composition (ratio of ¹³C/¹²C expressed as *δ*¹³C; Kelly 2000, Cherel & Hobson 2007) and their trophic levels at which the individuals are feeding through their nitrogen stable isotope composition (ratio of ¹⁵N/¹⁴N expressed as *δ*¹⁵N; McCutchan et al. 2003, Cherel & Hobson 2007). Stable isotope analyses can be performed on various types of biological tissues, notably on tissues that become inert, such as teeth and mystacial vibrissae (i.e., whiskers), providing dietary information spanning from several months to years (Hall-Aspland et al. 2005, Beltran et al. 2015, Jordaan et al. 2019). Inert tissues are usually sampled sequentially to assess chronological changes in stable isotope values providing insights into the long–term foraging habitats and trophic levels of individuals that travel in environments which are sometimes inaccessible to researchers (Hobson & Sease 1998, Authier et al. 2012, Lübcker et al. 2021). To correctly interpret chronological stable isotope patterns in phocid whiskers, it is essential to know the shedding dynamic and growth patterns of the whiskers as phocid species display contrasting patterns (reviewed in McHuron et al. 2020). Indeed, some species, such as spotted seals (*Phoca largha;* McHuron et al. 2016), ringed seals (*Pusa hispida;* McHuron et al. 2020) and southern elephant seals (*Mirounga leonina*; Lübcker et al. 2016), have considerable whisker loss during the annual fur moult. In contrast, other species show no apparent temporal pattern (e.g., leopard seals (*Hydrurga leptonyx*; Hall-Aspland et al. 2005), Hawaiian monk seals (*Neomonachus schauinslandi*) and bearded seals (*Erignathus barbatus*; McHuron et al. 2020). The whiskers of most of the phocid species follow an asymptotic growth pattern, characterised by an initial phase of linear growth that slows and ceases once the whisker approaches its asymptotical length (Greaves et al. 2004, Hall-Aspland et al. 2005, Lübcker et al. 2016, McHuron et al. 2016, 2020), typically conforming to a Von Bertalanffy growth curve (Von Bertalanffy 1938, 1957). The whisker remains on the muzzle until it sheds and is replaced by a new whisker. Conversely, in some phocid species, such as the bearded seal (*Erignathus barbatus*) and the Hawaiian monk seal (*Neomonachus schauinslandi*), whiskers grow continuously following a “rapid-slow” pattern until they are shed randomly throughout the year (McHuron et al. 2020). Despite the increasing application of stable isotope analyses of phocid whiskers to study seal foraging ecology and physiology (e.g. Hindell et al. 2012, Hückstädt et al. 2012, Baronia et al. 2025), information about growth and shedding dynamics are lacking for several species (McHuron et al. 2020). In Ross seals, the onset of whisker growth and the time period recorded within whiskers have never been investigated.

The multiple methods for defining whiskers shedding periods and recording time periods include measurements of regrowth rates of crimp beaded whiskers (Karpovich et al. 2022), photogrammetry (Greaves et al. 2004, Beltran et al. 2015, McHuron et al. 2016, 2020), the use of labelled stable isotopes (Hirons et al. 2001a, Zhao & Schell 2004) or the use of endogenous stable isotope variations along the whisker’s growing axis associated to known changes in diet, physiology or environment (Hall-Aspland et al. 2005, Lübcker et al. 2016, Rogers et al. 2016). Photogrammetry, recurrent monitoring of whisker growth or the use of labelled stable isotopes is impossible for Ross seals, because no individuals are currently kept under human care and recapturing specific individuals in the field is impractical. However, previous Ross seal tracking studies provided valuable information about their seasonal migrations and key periods of their annual biological cycle, such as nursing and moulting periods (Blix & Nordøy 2007, Arcalís-Planas et al. 2015, Wege et al. 2021, 2023). After completing their annual moult in January–February, lasting for about one week during which they are assumed to fast (Skinner & Klages 1994, Blix & Nordøy 2007), Ross seals travel north and remain in the open ocean south of the Antarctic Polar Front from March to October to forage (Blix & Nordøy 2007, Arcalís-Planas et al. 2015, Wege et al. 2021, 2023). They generally remain far from the ice edge during this open ocean period (Arcalís-Planas et al. 2015), but individuals have been observed to approach the ice edge, particularly around autumn (April to May; Wege et al. 2021). At the end of October, breeding females enter the consolidated pack–ice further south again to pup and nurse for approximately two weeks (Blix & Nordøy 2007, Wege et al. 2023), after which they travel north again in December to mate males (Thomas & Rogers 2009). To complete their biological cycle, Ross seals return to the pack ice to moult from January (Wege et al. 2021).

In the Southern Ocean, the carbon stable isotope values of particulate organic matter (POM) and primary producers at the base of the food web decrease with increasing latitude towards Antarctica (Francois et al. 1993, Espinasse et al. 2019, St John Glew et al. 2021). As carbon stable isotopes show little fractionation at each trophic step (generally between 0.4 – 0.8 ‰; Zanden & Rasmussen 2001, Post 2002), top predators have *δ*^13^C values close to the POM *δ*^13^C values of their foraging regions. Thus, *δ*^13^C values recorded in Ross seals’ whiskers would reflect the latitudinal *δ*^13^C POM variations during the seal’s south-north–south annual migrations, with higher *δ*^13^C values indicating foraging close to the Antarctic Polar Front, and lower *δ*^13^C values indicating foraging at higher latitudes close to the marginal ice zone. In the Indian sector of the Southern Ocean, Espinasse et al. (2019) reported a difference of 5 ‰ between the Antarctic Polar Front and the marginal ice zone, representing a rough linear gradient of approximately ∼0.3 ‰ per degree of latitude for POM. Given that the pack ice and the Polar Front are about 20° of latitude apart, an amplitude of *δ*¹³C variations on the order of 4-6 ‰ can be expected in Ross seal whiskers. Whilst the *δ*^13^C values of predators primarily reflect their foraging areas, their *δ*^15^N values provide information on their trophic level within an ecosystem (often also referred to as trophic position). Beyond the trophic information, tissues *δ*^15^N values are also affected by physiological processes such as pregnancy (Fuller et al. 2004, Kurle et al. 2014, Clark et al. 2016) and fasting (Hobson et al. 1993, Cherel et al. 2005). Recent meta-analyses demonstrated that fasting can cause a measurable increase in bulk tissue *δ*^15^N values up to 4 ‰ (Hertz et al. 2015, Doi et al. 2017). Therefore, we expected that Ross seals whisker *δ*^13^C profiles would reflect their latitudinal movements whereas higher *δ*^15^N values would relate to fasting during the moult period for males and females (early January – late February, ∼1 week, Blix & Nordøy 2007, Wege et al. 2023) and nursing period for females (late October – late November, ∼ 14 ± 2 d, Blix & Nordøy 2007, Wege et al. 2023). Slight variations in *δ*¹⁵N values, although smaller in magnitude, are also expected to reflect potential dietary changes among the different environments occupied (Skinner & Klages 1994, Constable et al. 2003, Blix & Nordøy 2007, Wege et al. 2021).

We hypothesised that the whiskers are shed annually during the fur moult and have a linear and continuous growth pattern until the whisker shed (**Figure 1A**). Thus, males are expected to show higher δ¹⁵N values near the whisker tip (i.e., distal end), reflecting the fasting phase occurring during the moult period (early January–late February), followed by variations along the whisker with lower δ¹⁵N values during oceanic foraging phases. Such a pattern would accentuate the variation already expected from the latitudinal baseline δ¹⁵N gradient in POM between marginal ice zone and the Antarctic Polar Front. For *δ*¹³C values, we expected more lower values at the whisker tip, reflecting foraging in high-latitude environments near Antarctica (St John Glew et al. 2021), followed by an increase to higher and then relatively constant *δ*¹³C values from March to December while males forage near the Antarctic Polar Front (Blix & Nordøy 2007, Arcalís-Planas et al. 2015, Wege et al. 2021). Within this generally steady pattern, decreases in *δ*¹³C are expected, likely reflecting a return to the pack ice in April–May for some individuals (Arcalís-Planas et al. 2015) and in December to moult. We also expected variations in *δ*^15^N values which would reflect the different proportion of fish, squid and krill to the diet of these seals (Skinner & Klages 1994), as prey availability varies between environments close to the pack ice and those in the open ocean (Xavier et al. 2006). Females are expected to show similar initial patterns as males (Blix & Nordøy 2007), with higher *δ*^15^N values and lower *δ*^13^C values at the whisker tip, followed by an increase in *δ*^13^C values corresponding to the northward migration (**Figure 1B**). For the females that pupped, we expected to see a decrease in *δ*^13^C values and a concurrent *δ*^15^N peak, representing the fasting associated with the nursing of the pup. If whisker growth is linear and constant, as in bearded and Hawaiian monk seals (McHuron et al. 2020), we should observe these variations in *δ*^13^C and *δ*^15^N values until the whisker shed. In the case of asymptotic growth — mainly observed in other phocids (Hirons et al. 2001b, Zhao & Schell 2004, Greaves et al. 2004, Beltran et al. 2015, Lübcker et al. 2016, McHuron et al. 2016, 2020) — we expected to see at least some stable isotope variations corresponding to the fasting linked to the annual fur moult and the northward migration occurring in late February - March.

**Figure 1:**
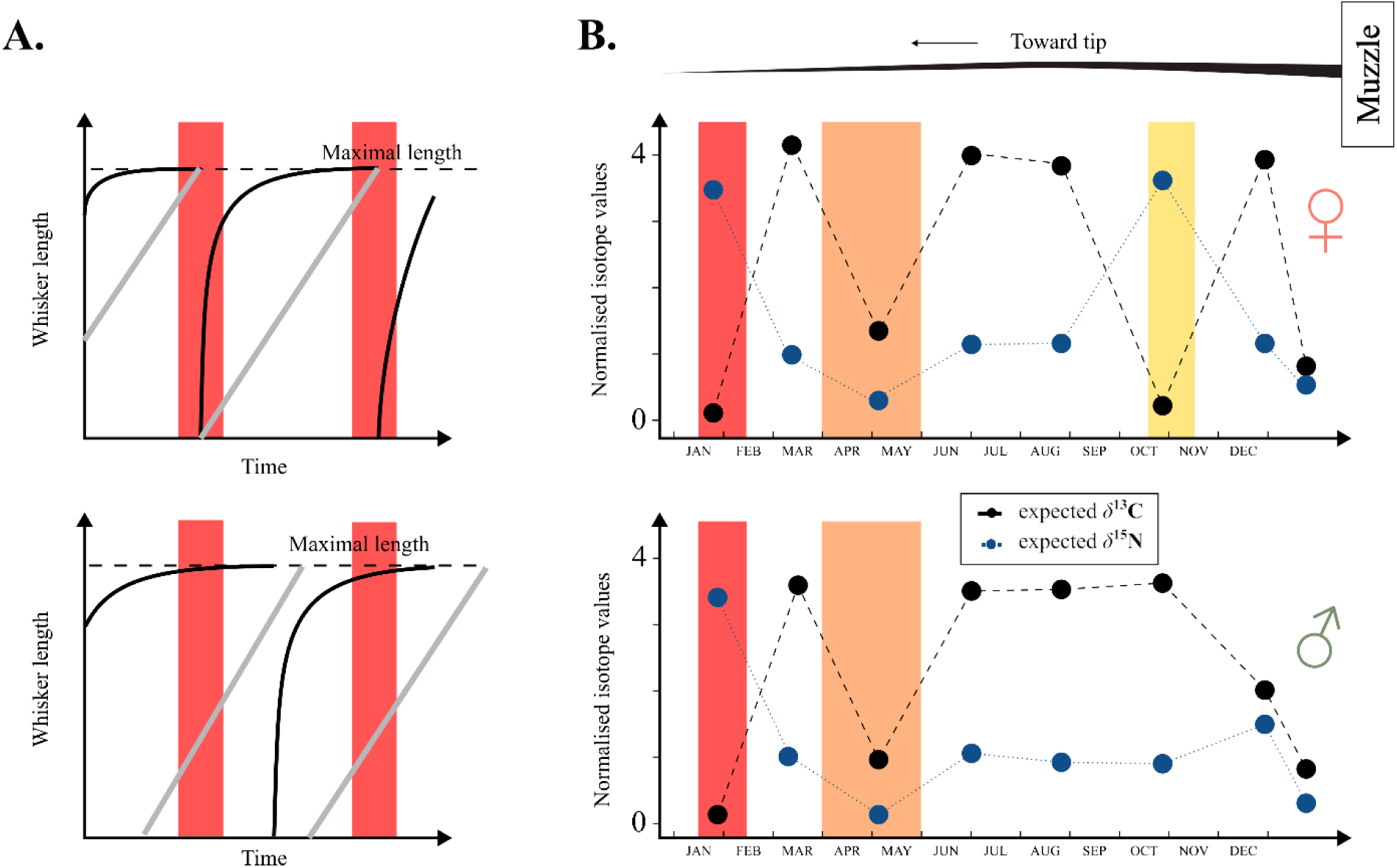
**A.** Schematic representation of an asymptotic (black) and a linear (grey) growth model regarding two temporal models of whisker loss (top: loss during the fur moulting period; bottom: loss of whiskers at random times). The red bar represents the moulting period. **B.** Pattern of expected variations in δ^13^C and δ^15^N values assuming a constant POM δ^15^N values, a very little influence of a potential change in diet, a whisker loss during the annual moulting period and whisker grow lasting for one year for females (top) and males (bottom). The red bar illustrates the moulting period, the orange one the period of return to the pack ice in April - May and the yellow one the breeding and nursing period in females.

Our study sought to achieve two objectives by using whisker *δ*^13^C and *δ*^15^N values and tracking data. The primary aim was to define the shedding phenology of Ross seals’ whiskers, with specific attention to ascertaining whether their shedding patterns coincide with the annual fur moult. Additionally, we attempted to determine the life-cycle period recorded in their whiskers, which is crucial information for future physiological and ecological studies that will be conducted using this tissue from this little-known species.

## 2. MATERIALS AND METHODS

### 2.1. Study area and sampled individuals

Twenty-one adult Ross seals were captured in the King Haakon VII Sea off Queen Maud Land (69°54′S–72°20′S, 2°00′W–17°46′W) during a research cruise aboard the *MV S.A. Agulhas II* in January 2016 (SANAE S55). Of these, 11 seals were fitted with satellite transmitters (see 2.2), and whisker samples (1–2 per individual) were collected from 14 seals (see 2.3). Six seals were both instrumented and sampled. In 2023, a further 28 adult Ross seals were captured in the eastern Weddell Sea off Queen Maud Land (70°51′S–69°93′S, 8°02′W–1°50′W) between January and February during the annual relief voyage to SANAE (Voyage 056) on the *MV S.A. Agulhas II*. Of these, 19 seals were equipped with satellite transmitters, and whisker samples (1–2 per individual) obtained from 26 seals. Seventeen seals were both instrumented and sampled. Detailed information on the sampled seals’ tagging/sampling date and location, standard length (cm, from snout to tail), and sex is provided in **Table 1**. Ross seals were studied and handled under the ethics approval of the University of Pretoria Animal Ethics Committee (EC082-15) and the University of South Africa College of Agriculture and Environmental Sciences (UNISA-CAES) Animal Research Ethics Committee (2021/CAES_AREC/089).

**Table 1:**
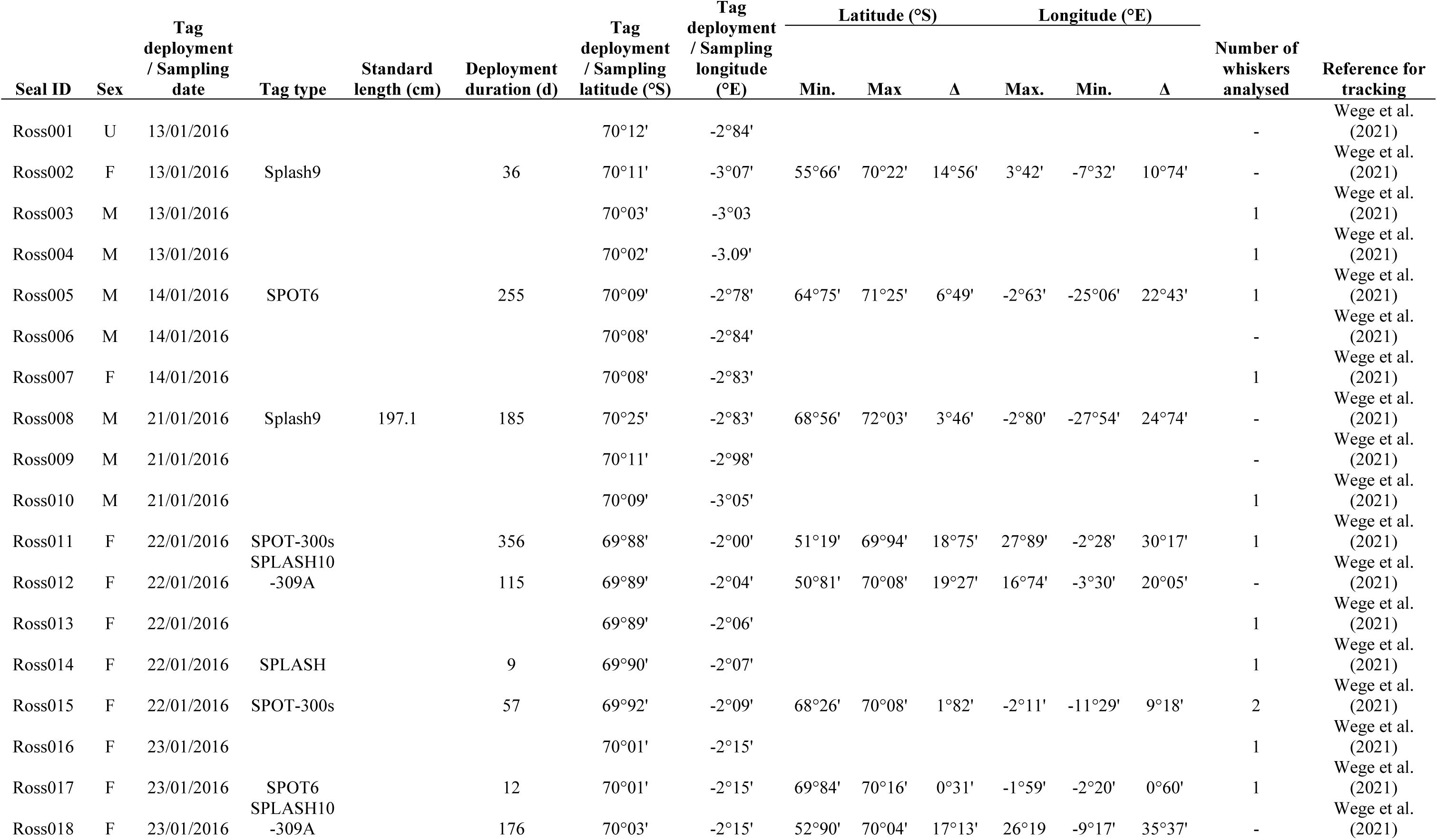
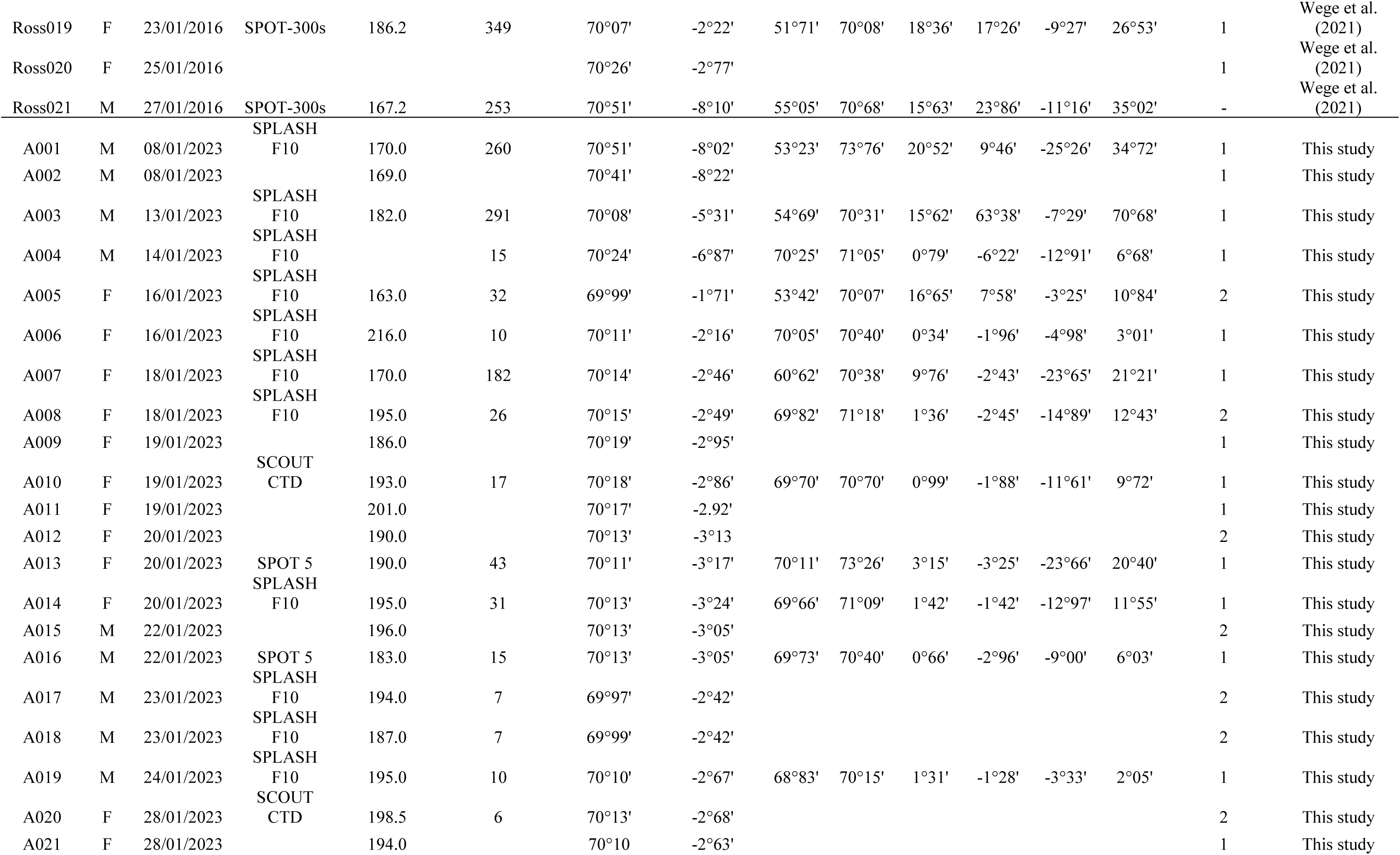
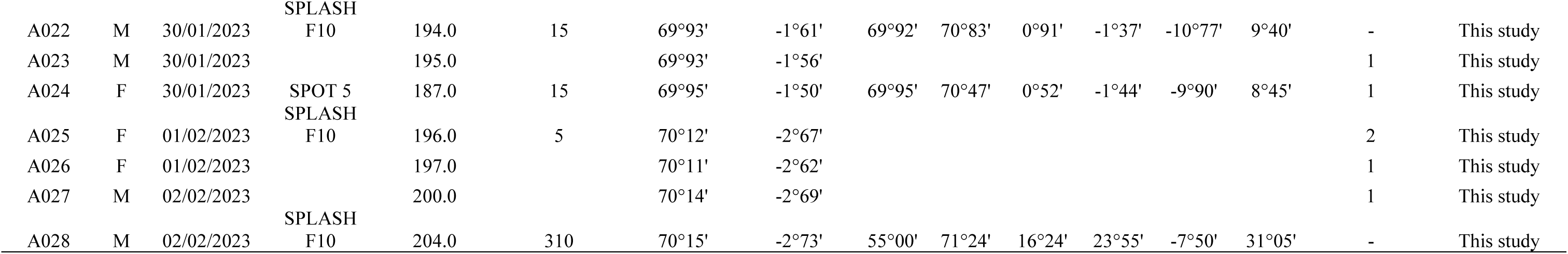
Biological data, date and location of tagging and sampling, and associated movements for Ross seals instrumented in 2016 and 2023.

### 2.2. Tag deployment and tracking data acquisition

The capture and tagging procedure followed Wege et al. (2021) where animals were handled for approximately 1 h. An A–frame net, comprising two stainless steel poles, hinged at the front end, with a nylon net in between, was used to capture the seals. After capturing the seal, a small hole was cut into the net where the top of the Ross seal’s head was located, to attach the satellite transmitter. Quicksetting Araldite Epoxy resin (AW2101/HW2951) or marine epoxy—Devcon 5 Minute Epoxy were used to glue the transmitter to the heads. Three types of satellite transmitters were used, SCOUT CTD, SPOT 5 and SPLASH F10 tags (Wildlife Computers, Redmond, WA, United States), all of which were Argos–linked (CLS, Toulouse, France; **Table 1**). We combine our Ross seals’ tracking data (2023) with published data (2016) collected in the same region during the South African National Antarctic Programme Expedition S55 (Wege et al. 2021).

The tracking data were filtered to exclude incomplete sequences and unrealistic point sequences, which are often encountered in ARGOS data. CLS Argos assigns a location class (LC = 3, 2, 1, 0, A, B or Z) to the calculated locations which indicates the accuracy of the location, with LC 3 being the most accurate. Vincent et al. (2002) suggested that the quality of LC = A or B may be acceptable and of interest in dealing with the distribution of seals that make large–scale movements within relatively short periods of time. Thus, locations from all classes were used in this study, except for those belonging to class Z, which are considered unusable. A speed–distance filter (Freitas 2012), setting a maximum swimming speed of 3 m.s^-1^, was applied (Arcalís-Planas et al. 2015), without turning angle filter. Interpolated data points were generated using a state–space model (Jonsen et al. 2013), implemented through the “*aniMotum*” R package (Jonsen et al. 2023). A 12-hour timestep was used as our aim was to investigate long-term movements and we only retained tracking records longer than 10 days (N = 22).

### 2.3. Whisker sampling, sample preparation and bulk carbon and nitrogen stable isotope composition measurements

One to two whiskers from 39 captured and sex-identified Ross seals (2016: N = 13 seals, 2023: N = 26 seals) were collected for bulk carbon and nitrogen stable isotope analyses (**Table 1**). The longest whisker located posterior closest to the eye, typically situated in the top or second-to-top row (Morgenthal et al. 2025), was selected for sampling. As the focus was placed on the longest whisker, the position on the muzzle of the sampled whisker varied, with no consistency observed either among individuals or between the left and right sides of the face when two whiskers were sampled. The whiskers were cut as close to the muzzle as possible because cortical cells keratinise at the base of the whiskers just above the bulb region of the follicle (Ling 1966), suggesting that the basal segment is still biologically active and does not represent pure keratin (Hückstädt et al. 2012, Lübcker et al. 2016). In cases, where whiskers were inadvertently plucked, the root portion (∼2 mm) was removed. The whiskers were placed in airtight bags and kept frozen at –20°C until they were prepared for stable isotope analysis.

Whiskers were first washed with deionized water and rinsed for 20 minutes with a mix of 1:2 chloroform: 70% ethanol solvent solution to remove surface contaminants. Samples were then rinsed twice with the chloroform: 70% ethanol solution and a final rinse with water. The whiskers were then dried for at least 24 h at 45°C. To obtain the target sample mass of 0.35–0.45 mg required for bulk stable isotope analysis, whiskers were subsampled in segments measuring 3.8 ± 3.3 mm (mean ± standard deviation, *n* = 436). The particularly small length of Ross seal whiskers (Morgenthal et al. 2025) necessitated the sampling of longer segments than those typically used in other phocid studies to obtain sufficient material for stable isotope analysis (Lübcker et al. 2020). The longest segments corresponded to the whisker tips, as the distal ends were thinner than the basal portions. Each segment was weighed into tin capsules (pre–cleaned in toluene) and the *δ*^13^C and *δ*^15^N values were measured on an Isolink CSNOH Flash elemental analyser or a Flash 2000 elemental analyser coupled to a Conflo IV gas control unit and linked to a Delta V plus continuous-flow isotope-ratio mass spectrometer (CF-IRMS, Thermo Fisher Scientific, Bremen, Germany) at Stable Light Isotope Laboratory, University of Cape Town (UCT), Cape Town, South Africa. A sub-set of 10 whiskers were analysed for *δ*^13^C and *δ*^15^N values using a Flash 1112 Series elemental analyser coupled to a Delta V Plus CF-IRMS (all Thermo Fisher Scientific, Bremen, Germany) in the Stable Isotope Laboratory of the Mammal Research Institute, University of Pretoria (UP), South Africa. Stable isotope results were calculated using Isodat© software (Thermo Fisher Scientific, Bremen, Germany) and are expressed as delta (*δ*) values in parts per thousand (per mil, ‰) relative to the international standard Vienna Pee Dee Belemnites (VPDB) for carbon and to atmospheric nitrogen (AIR) for nitrogen. To ensure the comparison of the results between the two analytical facilities, sample aliquots were interspersed with five internal standards (UCT and UP Merck gel, UCT and UP DL–Valine and UCT Sucrose) in each batch analysis to assess the analytical precision, which was ± 0.1 ‰ for both *δ*^13^C and *δ*^15^N values. These laboratory standards have been calibrated against international standards International Atomic Energy Agency (IAEA)-CH-3 (cellulose), IAEA-CH-6 (sucrose), IAEA-CH-7 (polyethylene foil), IAEA N-1 and IAEA N-2 (ammonium sulphate), IAEA NO-3 (potassium nitrate) in the Stable Isotope Laboratory of the Mammal Research Institute (UP) and with United States Geological Survey (USGS) 61 and USGS62 (both caffeine), casein from Elemental Microanalysis, USGS73 (L-valine), gelatin from Elemental MicroAnalysis and USGS66 (glycine) in the Stable Light Isotope Laboratory (UCT) in 2023. The atomic C:N ratios were also calculated.

### 2.4. Statistical analyses

Statistical analyses were done with R software, version 4.4 (R Core Team 2022). Statistical significance was set at p ≤ 0.05. We examined the effects of month, sex and their interaction on the spatial distribution (latitude and longitude visited by seals as response variables) using Generalised Additive Mixed Models (GAMMs) implemented via the *mgcv* R package (Wood 2017). The addition of an individual random effect accounted for the non-independence of repeated measurements within individuals and allowed the model to capture variability among individuals. Given that migrations are assumed to be annual, we grouped the sampling years together (2016 and 2023) to increase the statistical robustness of the GAMMs. To visually explore variations in *δ*¹³C and *δ*¹⁵N values along the length of the whiskers, we applied Locally Estimated Scatterplot Smoothing (LOESS) using the *ggplot2* R package (Wickham 2016). LOESS regression enabled us to visualise local trends in stable isotope values without assuming a predefined functional form, making it particularly appropriate for detecting non-linear patterns. To assess potential non-linear variations in stable isotope values along the whiskers, we used GAMMs. The distance from the tip, sex and sampling year were included as fixed effects, while whisker ID was incorporated as a random effect to account for repeated measurements within whiskers. Finally, to test whether stable isotope values differed at the tip compared with the rest of the whisker, we applied linear mixed-effects models (LMMs), with stable isotope values as the response, the position of the segment (“tip” arbitrarily defined as the last segment of the whisker vs. “rest” of the whisker), sex, and sampling year as fixed effects, and whisker ID as a random effect to account for repeated measurements within whiskers.

## 3. RESULTS

### 3.1. Tracking data

#### 3.1.1. Tag performances

Satellite tag performances are listed in **Table 1**. Of the 30 satellite tags deployed, five did not transmit a signal for more than 10 days and were removed from the dataset (2016: Ross014, 2023: A017, A018, A020 and A025). Five records from 2016 were also removed from the dataset because the tags failed (Ross005, Ross008, Ross014, Ross017; Wege et al. 2021). The mean recording duration for females equipped in 2016 was 182 days and ranged from 36 to 356 days (*n =* 6), and the 2023 mean recording duration was 44 days, ranging from 10 to 182 days (*n =* 8). The recording duration for the only male providing usable data (Ross021) in 2016 was 253 days. The mean recording duration for males equipped in 2023 was 131 days, ranging from 10 to 310 days (*n =* 7). A total of 40,288 locations (LC = B, or higher quality) were recorded, and each tag produced between 17 and 5,061 location estimates. Overall, 12% of all locations were of LC > 0, 2.4% of LC = 0, 10% of LC = A and 75% of LC = B.

#### 3.1.2. Ross seal movements

The state-space modelled location estimates for each instrumented Ross seal in 2016 and 2023 are represented in **Figure 2**. A summary of the latitudes and longitudes visited by Ross seal individuals is provided in **Table S1** while a second for each sex is provided **Table S2**.

**Figure 2:**
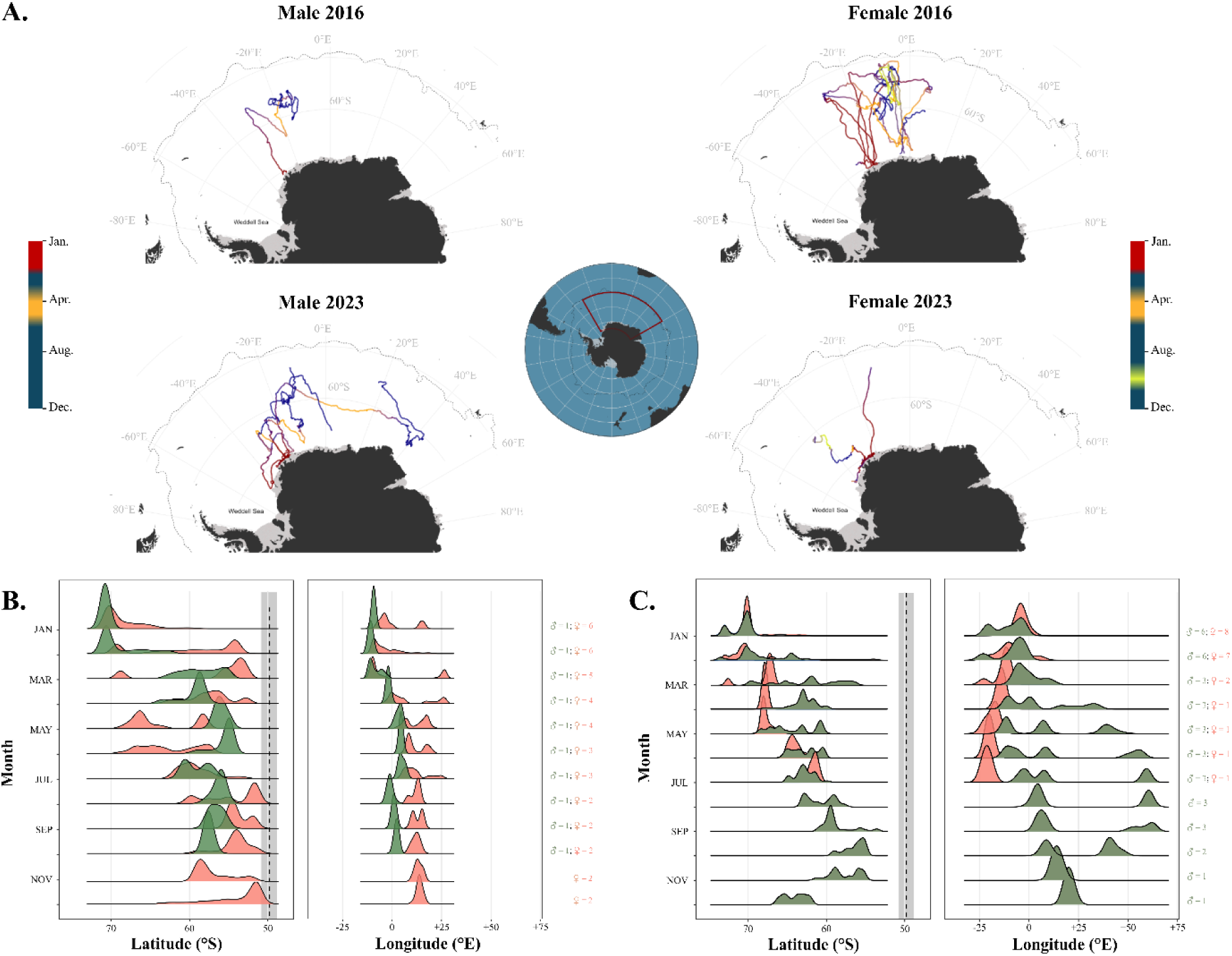
State–space modelled location estimates for instrumented Ross seal male (N= 1) and females (N= 6) in 2016 (Wege et al. 2021), as well as in males (N= 6) and females (N = 8) in 2023 (this study) in the eastern Weddell Sea off Queen Maud Land. The dashed line indicates the position of the Antarctic Polar Front (Orsi et al. 1995). Ice shelf extent is illustrated in grey area. Inset shows the location of the maps in relation to Antarctica, and the red rectangle outlines the study area. The colours of the tracks correspond to the different stages of Ross seal phenology: red for the moulting period, blue for the oceanic phases, orange for the return to the pack ice in April-May and yellow for the breeding period. Distribution of latitudes and longitudes visited by female (in red) and male (in green) Ross seals in 2016 (**B**.) and in 2023 (**C**.).

During the annual fur moult (January–February), both males and females remained in areas close to the pack ice between 69°S and 70°S (**Figure 2**). After moulting, individuals moved northward between March and April to areas south of the Antarctic Polar Front (67°S–55°S). The distance travelled during this period was farther in seals tracked in 2016 compared to those tracked in 2023, as in 2016 males and females were close to 55°S-60°S, whereas in 2023 they were between 60°S and 65°S. In 2016, a southward migration was observed from April to June, particularly in females. Then, from July to December, seals of both sexes remained near to and south of the Antarctic Polar Front. During December 2023, the last male that still transmitted data occupied environments at higher latitudes than those recorded between July and November. In terms of longitudinal distribution, males exhibited a much narrower range in 2016 compared to 2023, whereas females used a similar range of approximately 25° in both years (**Figure 2**). The wider distribution observed in males in 2023 likely results from the atypical movements of the seal A003 (**Table S1**).

The selected GAMM on latitudes visited by Ross seals involved the interaction between month and sex as fixed effect and explained 77.1% of the deviance (adj. R² = 0.77). It revealed a significant effect of month, with smooth terms for month differing between sexes (females: edf = 9.8, F = 19,184, p < 0.001; males: edf = 9.6, F = 20,544, p < 0.001, **Table 2**). The addition of an individual random effect (edf = 20.8, F = 302.6, p < 0.001) effectively accounted for inter–individual variability. The selected GAMM on longitudes visited by Ross seals involved the same fixed effect as the GAMM on latitudes. It explained 80.3% of the deviance (adj. R² = 0.80) and revealed a significant effect of month (females: edf = 8.1, F = 7,106, p < 0.001; males: edf = 9.7, F = 17,735, p < 0.001; **Table 2**). The inclusion of an individual random effect (edf = 44.5, F = 13.0, p < 0.001) captured substantial inter–individual variability.

**Table 2:** Results of generalised additive mixed models (GAMMs). Estimates, effective degrees of freedom (edf), standard errors (se), test statistics (t/F), p-values, and significance levels are reported. Significance codes: ns= non-significant, * = p < 0.05, **= p < 0.01, *** = p < 0.00. F = females, M = males, lat = latitude, lon = longitude.

| Term | Estimate/ edf | se | t/F value | p-value and significance level | Significance |
| --- | --- | --- | --- | --- | --- |
| **lat ~ year + s(month, sex) + s(1 \| seal ID)** |  |  |  |  |  |
| Intercept | -62.9 | 0.9 |  |  |  |
| s(month, F) | 9.8 |  | 19,184 | <0.001 | *** |
| s(month, M) | 9.6 |  | 20,544 | <0.001 | *** |
| s(1 \| seal ID) | 20.8 |  | 302.6 | <0.001 | *** |
| **lon ~ year + s(month, sex) + s(1 \| seal ID)** |  |  |  |  |  |
| Intercept | 5.3 | 2.3 |  |  |  |
| s(month, F) | 8.1 |  | 7,106 | <0.001 | *** |
| s(month, M) | 9.7 |  | 17,735 | <0.001 | *** |
| s(1 \| seal ID) | 20.8 |  | 622.3 | <0.001 | *** |
| **$\delta^{13}\text{C} \sim \text{year} + \text{s}(\text{dist. tip, sex}) + \text{s}(1 \vert \text{whisker ID})$** |  |  |  |  |  |
| Intercept (Year 2016) | -21.6 | 0.2 |  |  |  |
| Year 2023 | -1.0 | 0.3 | -3.3 | 0.001 | ** |
| s(dist. tip, F) | 4.3 |  | 2.8 | 0.014 | * |
| s(dist. tip, M) | 1.8 |  | 3.1 | 0.028 | * |
| s(1 \| whisker ID) | 44.7 |  | 40.6 | <0.001 | *** |
| **$\delta^{15}\text{N} \sim \text{year} + \text{s}(\text{dist. tip, sex}) + \text{s}(1 \vert \text{whisker ID})$** |  |  |  |  |  |
| Intercept (Year 2016) | 9.8 | 0.1 |  |  |  |
| Year 2023 | -0.2 | 0.2 | -1.1 | 0.282 | ns |
| s(dist. tip, F) | 5.0 |  | 9.2 | <0.001 | *** |
| s(dist. tip, M) | 2.7 |  | 5.3 | <0.001 | *** |
| s(1 \| whisker ID) | 44.5 |  | 13.0 | <0.001 | *** |

### 3.2. Ross seal whisker *δ*^13^C and *δ*^15^N values

The total length of sampled Ross seal whiskers was between 18 and 52 mm (35.4 ± 8.2 mm, *n =* 48). Each whisker was cut into 5 to 17 segments with subsampled length ranging from 1 to 20 mm, and an overall mean subsampled length of 3.8 ± 3.3 mm. Carbon and nitrogen stable isotopes values are reported in **Table S3**. The *δ*¹³C values measured in all Ross seal whiskers ranged from –24.8 to –19.6 ‰. Within individual whiskers, *δ*¹³C values varied between 0.3 and 3.2 ‰, with a mean value of 1.2 ± 0.7 ‰. The *δ*¹⁵N values of all whiskers ranged from 7.9 to 11.7 ‰, with individual intra–whisker *δ*¹⁵N values ranging from 0.4 to 2.1 ‰ and a mean of 1.1 ± 0.5 ‰ (**Table 3**).

**Table 3:** Summary of δ¹³C and δ¹⁵N values measured in whisker segments of Ross seals, presented by sex (F = female, M = male) and year. N refers to the number of individuals, the number in bracket the number of whiskers analysed, and n to the number of segments analysed.

| Year | Sex | N | n | $\delta^{13}\text{C}$ (‰, VPDB) | | | $\delta^{15}\text{N}$ (‰, AIR) | | |
| --- | --- | --- | --- | --- | --- | --- | --- | --- | --- |
| | | | | Range (‰) | Mean range $\pm 1$ SD | Intra-whisker variability (‰) | Range (‰) | Mean range $\pm 1$ SD | Intra-whisker variability (‰) |
| All | F | 24 | 260 | -24.8 to -19.6 | $1.2 \pm 0.7$ | 0.3 – 3.2 | 7.9 to 11.3 | $1.1 \pm 0.5$ | 0.4 – 2.0 |
| | M | 15 | 176 | -23.9 to -20.2 | $1.2 \pm 0.6$ | 0.2 – 2.2 | 8.2 to 11.7 | $1.1 \pm 0.5$ | 0.4 – 2.1 |
| 2016 | F | 9 (10) | 90 | -23.6 to -19.6 | $1.0 \pm 0.6$ | 0.3 – 2.1 | 8.9 to 11.3 | $1.2 \pm 0.6$ | 0.4 – 2.0 |
| | M | 4 (4) | 43 | -23.3 to -20.6 | $1.1 \pm 0.5$ | 0.2 – 1.8 | 9.8 to 11.1 | $1.0 \pm 0.5$ | 0.8 – 1.8 |
| 2023 | F | 15 (20) | 170 | -24.8 to -20.4 | $1.4 \pm 0.7$ | 0.7 – 3.1 | 7.9 to 11.0 | $1.1 \pm 0.4$ | 0.5 – 2.2 |
| | M | 11 (14) | 133 | -24.0 to -20.4 | $1.1 \pm 0.5$ | 0.4 – 2.2 | 8.3 to 11.7 | $1.0 \pm 0.5$ | 0.4 – 2.1 |

GAMMs on Ross seals geographical distribution indicated a strong influence of month on the latitudes and longitudes visited by the seals. Therefore, based on our working hypothesis regarding migrations and the latitudinal gradient of POM *δ*¹³C values, we expected that the variations in *δ*¹³C values along the whiskers (representing temporal progression) would be influenced by the distance to the whisker tip. The selected GAMM for *δ*¹³C involving the sampling year and the interaction between the distance from the tip with sex (**Table 2**), explained 85% of the variance in *δ*¹³C values (adj. R² = 0.83). The model indicated that values from whiskers sampled in 2023 are significantly lower by 1 ‰ than those sampled in 2016 (t = -3.3, p < 0.001). It also indicated that *δ*¹³C values showed a significant non-linear relationship with the distance from the tip in males’ whiskers (edf = 1.8, F = 3.1, p = 0.028) and in females’ whiskers (edf = 4.3, F = 2.8, p = 0.014). Random effects of individual samples accounted for a substantial portion of the variance (edf = 44.7, F = 40.6, p < 0.001). The *δ*¹⁵N GAMM, including the interaction between the distance from the whisker tip and sex and year as fixed effects, explained 68.2% of the variance in *δ*¹⁵N values (adj. R² = 0.64). The *δ*¹⁵N values were not significantly different between whiskers sampled in 2016 and 2023 (p = 0.282, **Table 2**). A significant non-linear relationship was evident between *δ*¹⁵N values and distance from the whisker tip with sex (F: edf = 5.0, F = 9.2, p < 0.001, M: edf = 2.7, F = 5.3, p < 0.001). Random effects of individual samples accounted for a large portion of the variance (edf = 44.5, F = 12.9, p < 0.001).

The *δ*¹³C and *δ*¹⁵N profiles of the whiskers did not match our working hypothesis, as no similar or aligned stable isotope profiles were observed between sexes and seals of the same sex (see **Figure 3** and **Figure S1** for individual results).

**Figure 3:**
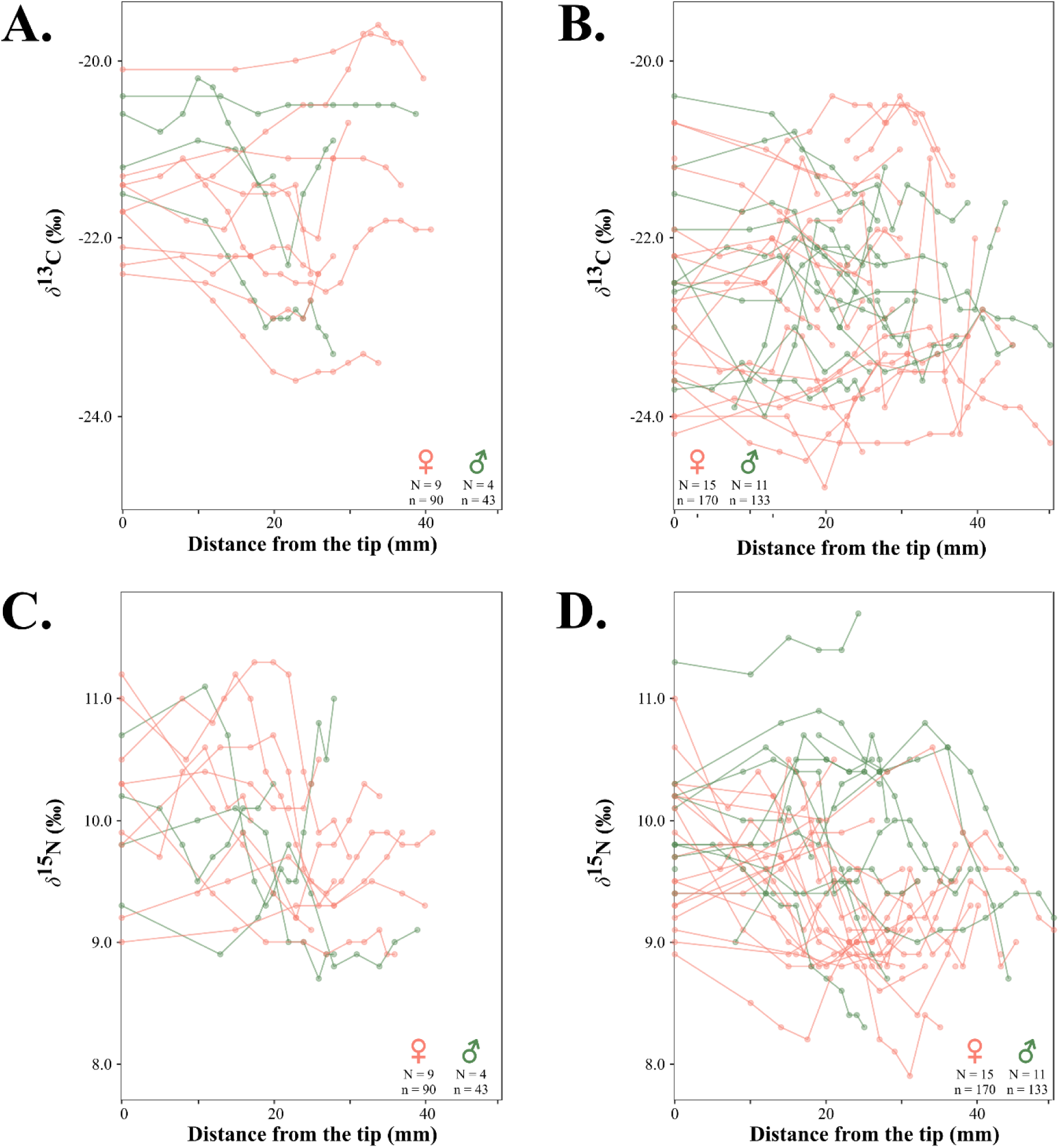
Whisker δ^13^C values along the whisker length according to sex (red for females and green for males) for seals sampled in 2016 (**A.**) and in 2023 (**B.).** Whisker δ^15^N values along the whisker length according to sex for seals sampled in 2016 (**C.**) and in 2023 (**D.).** Each straight-line profile represents one whisker. N corresponds to the number of seals and n, the number of whisker segments analysed.

The distribution of *δ*¹³C and *δ*¹⁵N values did not systematically indicate lower *δ*¹³C values at the tips, nor higher *δ*¹⁵N values (**Figure 4**). To formally assess this absence of observed stable isotope composition differences between the tip and the rest of the whisker, we fitted LMMs including position of the segment along the whisker (tip vs. rest of the whisker), sex, and year as fixed effects, with whisker identity as a random intercept. For *δ*¹³C, the model including position, sex, and year as fixed effects, with whisker identity as a random intercept, indicated an intercept of –21.6 ± 0.3 ‰ and neither the distance from the tip nor sex had significant effects, whereas year had a significant negative effect (**Table 4**). Random intercept variance was 0.80 (SD = 0.90), with residual variance of 0.22 (SD = 0.47), indicating inter-individual differences in *δ*¹³C values. Overall, *δ*¹³C values appeared largely independent of the position along the whisker and sex but decreased in 2023. For δ¹⁵N values, the model including segment position, sex and year as fixed effects, with individual identity as a random intercept, revealed an intercept of 9.5 ± 0.1 ‰. The sex as well as the distance from the tip had significant positive effect, with higher *δ*¹⁵N values in tips in male whiskers (0.4 ± 0.2 ‰). However, the mean *δ*¹⁵N difference between tips and the rest of the whiskers was relatively small (0.3 ± 0.1 ‰). The sampling year had no significant effect on whiskers’ *δ*¹⁵N values (**Table 4**). Random intercept variance was 0.23 (SD = 0.48), with residual variance of 0.17 (SD = 0.42), reflecting moderate inter-individual variability but a clear within-whisker effect of position.

**Figure 4:**
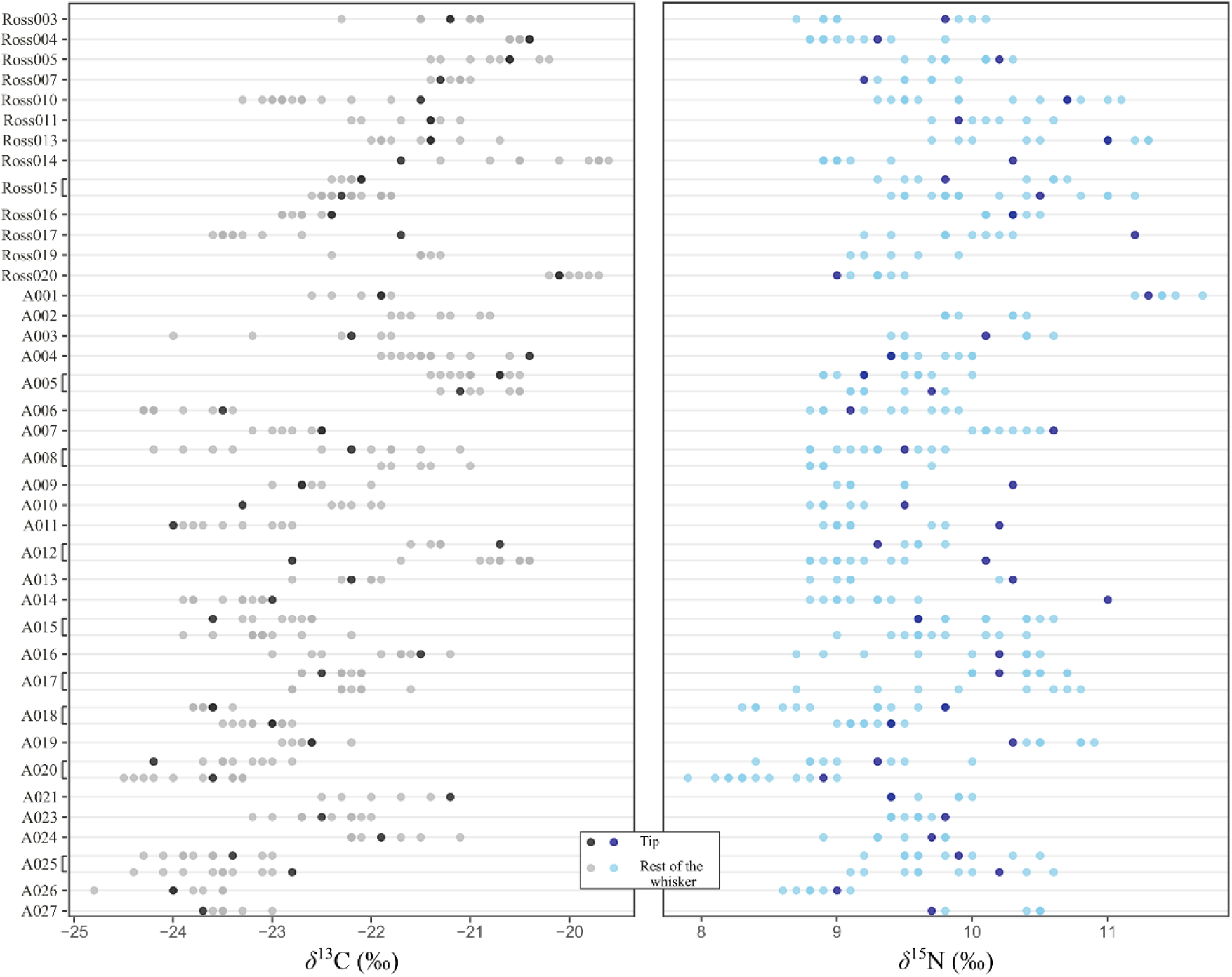
Distribution of δ¹^3^C and δ¹⁵N values per whisker, with comparison between tip and the rest of the whisker.

**Table 4:** Results of linear mixed models (LMMs) testing the effects of the segment position (tip vs. rest of the whisker), sex, and year on δ¹³C and δ¹⁵N values. Estimates are presented with standard errors (SE), degrees of freedom (df), test statistics (t-values), p-values and 95% confidence intervals (CI). Significance levels are reported by asterisks (*: p < 0.05, ***: p < 0.001). The reference year is 2016 and the reference sex is the female sex.

| Response variable | Predictor | Estimate ± SE | df | t-value | p-value | 95% CI |
| --- | --- | --- | --- | --- | --- | --- |
| $\delta^{13}\text{C}$ | Intercept | -21.6 ± 0.3 | | | | [-22.1, -21.1] |
|  | Segment position | 0.1 ± 0.1 | 374 | 1 | 0.4 | [-0.07, 0.2] |
|  | Sex (M) | 0.1 ± 0.3 | 45 | 0.4 | 0.7 | [-0.4, 0.7] |
|  | Year (2023) | -1.0 ± 0.3 | 45.0 | -3.3 | 0.002 | [-1.6, -0.4]** |
| $\delta^{15}\text{N}$ | Intercept | 9.5 ± 0.1 | | | | [9.3, 9.7] |
|  | Segment position | 0.3 ± 0.1 | 375 | 4.1 | <0.001 | [0.1, 0.4]*** |
|  | Sex (M) | 0.4 ± 0.2 | 46 | 2.5 | 0.02 | [0.1, 0.7]* |

Finally, the comparison of *δ*¹³C and *δ*¹⁵N values between one left and one right whisker of the same individual revealed variable stable isotope profiles (**Figure 5**). In some seals (e.g. A012, A017), no similarity was observed between the stable isotope profiles of the two whiskers for either *δ*¹³C or *δ*¹⁵N values, whereas in other seals, patterns of stable isotope variations were similar but never perfectly aligned or superimposable (e.g. A008, A015, A025). Interestingly, when similar trends were observed for *δ*¹³C values, a comparable pattern was generally seen for *δ*¹⁵N values, indicating a coordinated variation in *δ*¹³C and *δ*¹⁵N values within individuals.

**Figure 5:**
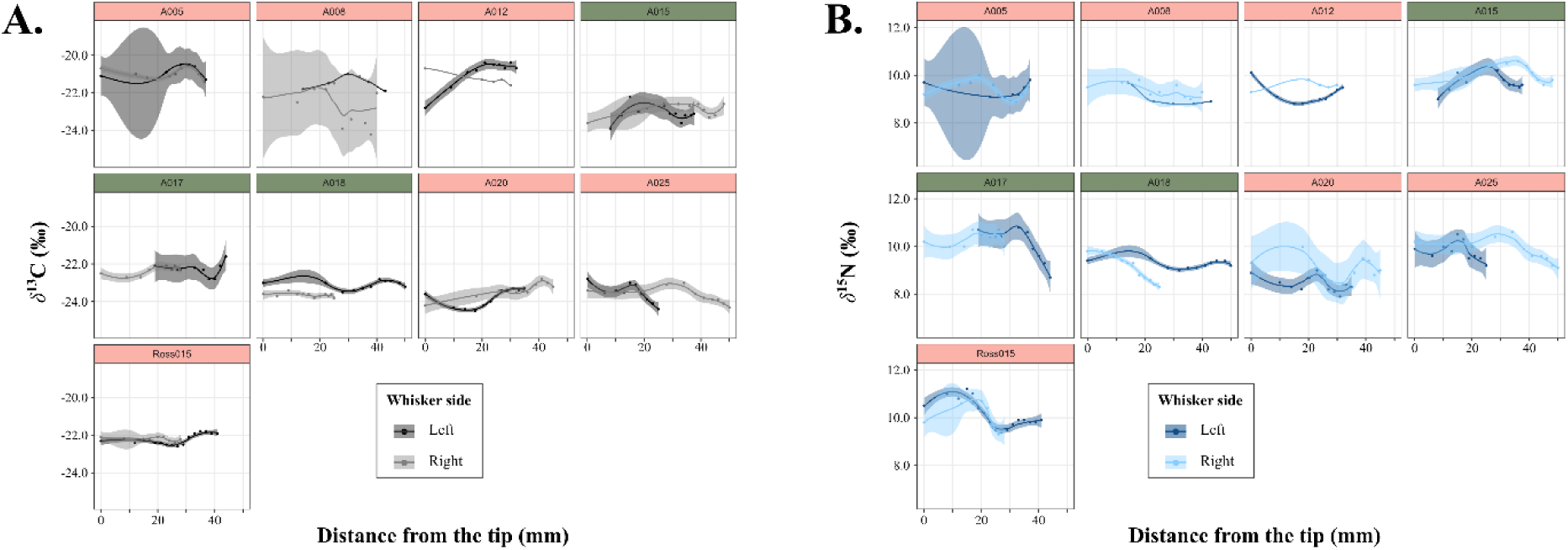
Comparison of **A.** δ¹³C and **B.** δ¹⁵N values along the left and right whiskers of individual Ross seals. The colour of the panels indicates the sex of the individual (green = male, red = female). The curves represent the local regression (LOESS) with their associated 95% confidence intervals.

## 4. DISCUSSION

Our study aimed to determine whether Ross seal whiskers’ *δ*¹³C and *δ*¹^5^N profiles, supplemented with tracking data collected after whisker growth, could provide information on the period during which the whiskers shed, and the period captured by the whiskers. We suggest that Ross seal whiskers are not shed preferentially in the fasting period occurrence during the annual fur moult. Moreover, given our inability to identify the key life events of Ross seals in their whiskers, mean whisker growth rates and the temporal window represented by the tissue cannot be precisely estimated. Our difficulties in identifying fasting events and seasonal migrations likely reflect a combination of environmental factors (spatial and temporal variability of POM *δ*¹³C values in the Southern Ocean), factors intrinsic to the physiology of Ross seals such as non-seasonal shedding, growth rate of the whiskers, and the insufficient stable isotopic resolution of whisker records relative to the timing of fasting periods. The rationale behind these finding are discussed in more detail below.

### 4.1. Spatial and temporal variability of particulate organic matter *δ*¹³C values in the Southern Ocean: a challenge for interpreting Ross seal latitudinal migrations from their whiskers?

Latitudinal gradients in the *δ*¹³C values of surface POM have been well documented in the Southern Ocean (e.g., Francois et al. 1993, Goericke & Fry 1994, Espinasse et al. 2019, St John Glew et al. 2021). The decrease in *δ*¹³C values associated with increasing latitude primarily results from the availability of dissolved CO_2_ which is largely influenced by seawater temperature (Rau et al. 1991, Goericke & Fry 1994) and thus latitude in the Southern Ocean (Stark et al. 2019). However, additional factors, such as time (season or year as shown in our *δ*¹³C GAMM), continent proximity, the intensity of primary production, variations in phytoplankton community composition, growth phase, productivity and cell size (Goericke & Fry 1994, Laws et al. 1997, Kukert & Riebesell 1998, Riebesell et al. 2000a b), may also contribute to POM *δ*¹³C variability. In the marginal ice zone where there is ice melt for example, primary productivity is increased through enhanced iron input, producing higher *δ*¹³C values than those predicted by the latitudinal gradient (Munro et al. 2010, Bury et al. 2024). POM *δ*¹³C values do not vary linearly with latitude but have abrupt transitions at oceanic fronts (Francois et al. 1993, Cherel & Hobson 2007). This, can present challenges in detecting latitudinal animal migrations using incremental *δ*¹³C variations along Ross seal whiskers as they remain south of the Antarctic Polar Front (Blix & Nordøy 2007, Arcalís-Planas et al. 2015, Wege et al. 2021, this study).

In addition, benthic environments possess higher POM *δ*¹³C values compared to pelagic and offshore environments (France 1995, Hobson et al. 1995), a factor that could lead to misinterpretations of the whisker *δ*¹³C records if Ross seals were consuming benthic prey at particular time of the year. Stomach content analyses revealed that both males and females Ross seals prey on fish (*Pleurogramma antarctica*), squid (*Psychroteuthis glacialis, Alluroteuthis antarcticus*) and krill (*Euphausia superba*) (Øritsland 1977, Laws 1984, Rau et al. 1992, Skinner & Klages 1994). All these preys species live at depths between 50 and 300 m (i.e. in the epipelagic zone) over deep ocean basins (Bengtson & Stewart 1997, Blix & Nordøy 2007, Wege et al. 2023). Although Ross seals are capable of benthic dives when they occupy the marginal ice zone in late January – late February, stomach content information as well as new diving data records (only 0.07% of the dives recorded were close to the continental shelf; Lewis, unpublished data) does not indicate extensive benthic foraging.

### 4.2. Ross seal whiskers shedding phenology

#### 4.2.1. No preferential whisker shedding during the fur moulting period

The *δ*¹⁵N LMM showed that whisker tips have significantly higher *δ*¹⁵N values compared to the rest of the whisker; however, the small magnitude of this difference (+ 0.3 ‰) does not provide sufficient evidence to clearly attribute it to fasting during the annual fur moulting although the *δ*¹⁵N peak could be diluted because of the length of whiskers’ tip segments. On the other hand, no significant differences were observed for *δ*¹³C values related to the segment position. The only observed *δ*¹³C value difference was between sampling years which are more likely associated with environmental changes that occurred between 2016 and 2023, potentially as a consequence of global warming and changes in hydrographic and biological conditions (Mestre et al. 2020, Séon et al. 2025). Moreover, the *δ*¹³C and *δ*¹^5^N profiles along Ross seal whiskers did not reveal a consistent pattern of higher *δ*¹⁵N values or lower *δ*¹³C values at the tips or any consistent stable isotopic patterns that would be expected if whisker replacement happened at the same time as the annual moult of the fur. This contrasts with our working hypothesis that whiskers are preferentially shed during the fasting period associated with the annual fur moult, as has been reported in some other phocid species (Beltran et al. 2015, Lübcker et al. 2016, McHuron et al. 2020). However, this corresponds to the absence of any trend observed in the number of whiskers still present on the muzzle at the time of sampling for seals at different stages of moulting. The aseasonal whisker shedding is not an exception and is observed in several other phocid species such as gray seals (Greaves et al. 2004), monk seals, bearded seals (McHuron et al. 2020), leopard seals (Hall-Aspland et al. 2005) and harbour seals (Hirons et al. 2001, Zhao & Schell 2004, Karpovich et al. 2022). Higher *δ*¹⁵N values and lower *δ*¹³C values were observed at the tips of the whiskers in some individuals (A003, A009, A011, A014; 1 male and 3 females). However, this pattern was observed in only a small proportion of the whiskers sampled (8%) and cannot be considered a general trend.

In a case where whiskers are lost and replaced simultaneously, they should exhibit a similar stable isotopic profile along their entire length, assuming that the growth rate is identical between whiskers (e.g. Lewis et al. 2006). In Ross seals, carbon and nitrogen stable isotope profiles obtained from two whiskers of the same seal displayed neither similar stable isotope variations, nor temporal alignment. In three of the nine seals sampled for both left and right whiskers (A008, A015, A025), we observed similar patterns of variation in *δ*¹³C and *δ*¹⁵N values between the two whiskers. However, these patterns were offset relative to the distance from the whisker tip. This misalignment suggests that despite both whiskers recording comparable ecological information, they were not shed and regrown synchronously, providing additional evidence that whisker replacement does not occur synchronously or seasonally.

#### 4.2.2. Limited interpretations of fasting periods from *δ*¹⁵N values

Ross seals experience one to two distinct, temporally constrained periods within their annual biological cycle during which they fast or significantly reduce their food intake (moulting period for both sexes and the breeding period for females). As mentioned above, although higher *δ*¹⁵N values were detected at the whisker tip, the difference (0.3 ± 0.1 ‰) is too small to be attributed to fasting. Several factors may contribute to explain our inability to clearly identify a clear *δ*¹⁵N peak corresponding to fasting. Firstly, whilst Ross seals are assumed to fast during the moulting period (Skinner & Klages 1994), they continue to infrequently dive (Blix & Nordøy 2007) indicative of a substantial persistence of foraging behaviour and this regardless of the stage of moulting (Lewis, unpublished data). This residual foraging could mitigate the effect of fasting on *δ*¹⁵N values recorded in whiskers, since the duration of fasting is a key parameter for the increase in *δ*¹⁵N values to be significant (Hertz et al. 2015). Moreover, the relatively brief duration of the moulting phase (∼ 1 week, Blix & Nordøy (2007) compared to southern elephant seals for example: 30.1 ± 9.7 days, Lübcker et al. 2016), combined with the unknown growth rate of the whiskers, may further reduce the resolution with which stable isotope variations can be detected.

The breeding season is characterised by considerable behavioural variability among Ross seal females. Recent biologging data suggest that females lactate for approximately two weeks (Blix & Nordøy 2007, Wege et al. 2023), during which, some females remain hauled out continuously for periods of up to 7–9 days, whilst others alternate between haul-out intervals and aquatic activity (Wege et al. 2023). Arcalís–Planas et al. (2015) and Wege et al. (2023) highlighted significant inter-individual differences in haul-out durations during the breeding season, which could reflect the coexistence of flexible reproductive strategies in Ross seals similar to those found in Weddell seals (Wheatley et al. 2008) and harbour seals (Bowen et al. 2001). Specifically, some females could be capital breeders, relying on stored energy reserves, whereas others could be facultative income breeders in some years, intermittently engaging in foraging during pup-rearing. This variability in fasting behaviour among individuals could explain the difficulty of identifying marked *δ*¹⁵N peaks attributable to fasting periods compared to other species such as southern elephant seals that are strict capital breeders (Lübcker et al. 2020).

Furthermore, it is also possible that the expected increase in *δ*¹⁵N values in the whiskers during fasting periods (Hobson et al. 1993, Iverson et al. 1995) may be absent because energy is allocated to other physiological processes instead of whisker synthesis (Hilderbrand et al. 1996). Indeed, moulting and nursing require a lot of energy from the female (Champagne et al. 2012), so it is conceivable that the synthesis of whisker tissue does not occur during the moulting period in order to conserve energy and maintain functional whiskers (Hirons et al. 2001b, Greaves et al. 2004). The Ross seal is one of the smallest phocid species (∼ 200 kg, Laws 1984). Thus, the energy allocated to physiological processes may be a limiting factor, unlike larger species such as the southern elephant seal, whose mass (> 400 kg) may make it possible to fast during moulting and to synthesise new functional whiskers rapidly. It is also important to note that, although we have taken precautions by selecting the least damaged whiskers, some vibrissae may have damaged ends, which would cause the early stages of whisker growth to disappear over time due to the significant mechanical forces exerted on them (Morgenthal et al. 2025).

Finally, we must report some technical limitations. To have the minimum sample mass to carry out the stable isotope analyses, we sometimes had to use relatively long whisker segments (i.e. at the tip up to 20 mm due to the narrower diameter). This could obscure the detection of small-scale temporal variations in diet or physiological state, reducing the ability to resolve short-term life-history events through stable isotope analysis. For Ross seals, this is potentially a major methodological constraint given the small size of their whiskers (Polkey & Bonner 1966) compared to other phocid species (e.g. McHuron et al. 2020). With technical advances in instrumentation and the ability to accurately analyse smaller quantities of whisker material, future analytical capabilities may open up opportunities to provide greater sampling resolution to further investigate Ross seal ecology and phenology.

### 4.3 Hypothesis regarding Ross seal whiskers shedding and growth dynamic

In the absence of species-specific whisker growth data, McHuron et al. (2020) suggested extrapolating whisker growth rates from phylogenetically and ecologically similar species, although precautions should be applied in light of the ecological diversity of phocids. This approach needs to consider factors such as comparable body size, feeding ecology, whisker morphology, and phylogenetic proximity. From a phylogenetic perspective, the Ross seal, as Lobodontini, is closely related to the crabeater seal (*Lobodon carcinophaga*), the Weddell seal (*Leptonychotes weddellii*) and the leopard seal (*Hydrurga leptonyx*, Berta et al. 2018). Currently, data on whisker growth rate for crabeater seals are lacking, but available for Weddell seal pups only (Baronia et al. 2025), which likely differ in growth patterns from adults. Applying whisker growth rates from leopard seals that have an average whisker growth rate of 0.2 mm.d^-1^, suggests that for a maximal whisker length of 5.2 cm in the Ross seal, the period captured could represent up to 260 days. Alternatively, when considering the growth rate range reported across other phocid seals (0.3 mm.d^-1^ to 0.9 mm.d^-1^; McHuron et al., 2020), the estimated period captured by Ross seal total whisker lengths would range from 60 to 170 days.

## 5. CONCLUSION

Our new tracking data confirmed the known annual latitudinal migrations of Ross seals. Based on the established latitudinal gradients of *δ*¹³C in POM in the Southern Ocean, as well as the influence of fasting periods on *δ*¹⁵N values in biological tissues, the analysis of *δ*¹³C and *δ*¹⁵N values along the whiskers of both adult males and females allowed us to investigate the shedding dynamic and whisker growth patterns in the Ross seal. We suggest no preferential whisker shedding during the fur moulting period as incremental *δ*¹⁵N values along the whiskers did not allow us to clearly identify fasting periods related to moulting or breeding periods. Our study contributes new data on stable isotope whisker composition and shedding patterns for the Monachinae subfamily. By providing novel biological information, our study highlights the periods of the life cycle that these tissues can record. In particular, it offers a valuable foundation for future research on the Ross seal and holds the potential to generate critical insights into this animal’s ecological role and trophic dynamics.

## ACKNOWLEDGMENTS

The authors acknowledge the UCT Stable Light Isotope Laboratory and the Biogeochemistry Research Infrastructure Platform (BIOGRIP, University of Cape Town, South Africa) and Stable Isotope Laboratory of the Mammal Research Institute (University of Pretoria, South Africa) for performing the stable isotope measurements. This work is supported by the National Research Foundation of South Africa (SANAP200317509817). Dr. Martin Postma, Dr. Nico Lübcker, Wiam Haddad and Andre van Tonder are thanked for their assistance in the field. The Officers and Crew of the MV SA Agulhas II and RV Polarstern extended every possible courtesy to us in support of our research objectives, especially Captain Gavin Syndercombe. Chief Scientists of the respective voyages, Dr Thato Mtshali, Dr Mike Schröder, Dr Tommy Ryan-Keogh and Gavin Louw are thanked for their support and the Department of Environment Affairs’ Co-ordinating Officer (DCO) and Deputy DCO, for facilitation.

---

**Figure S1:**
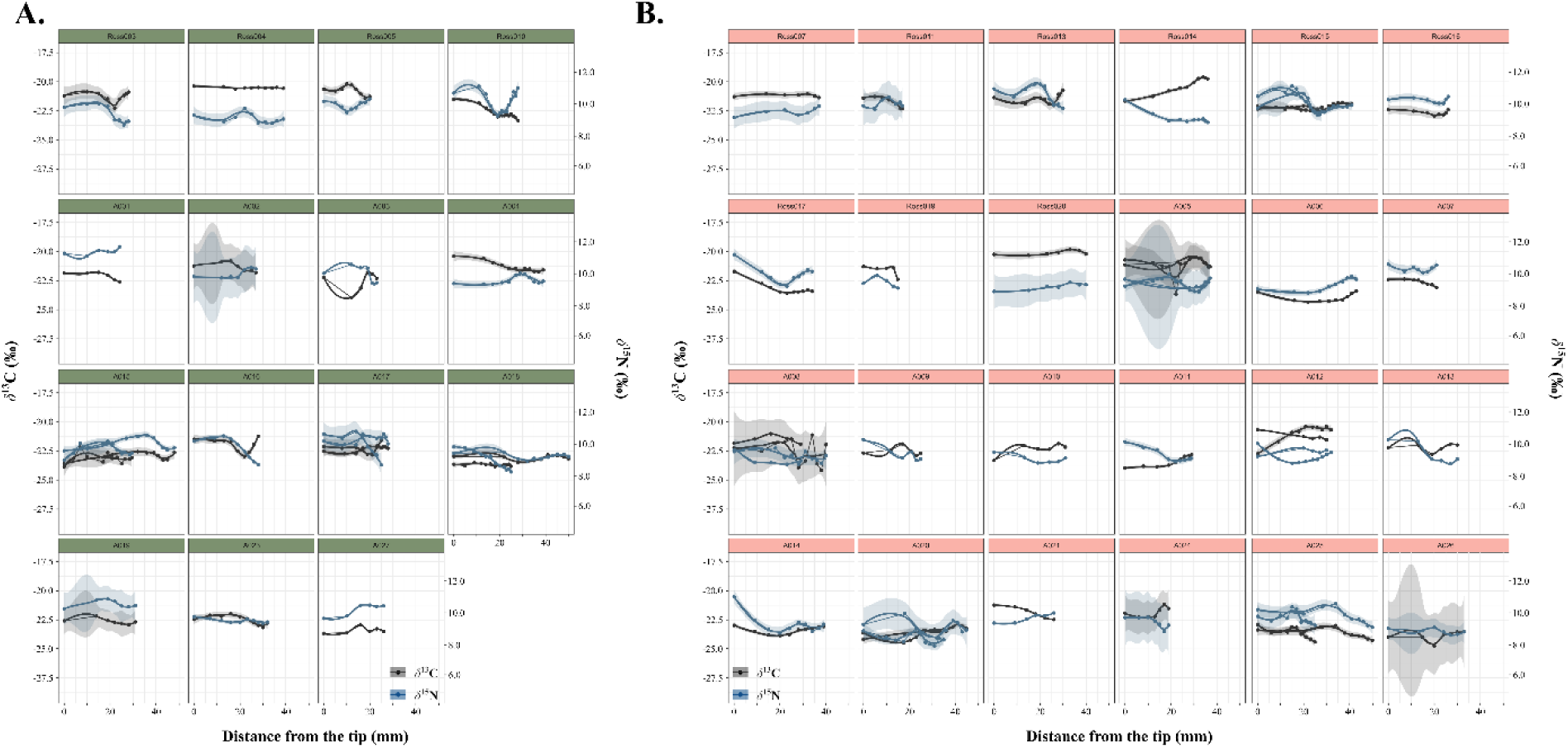
Carbon and nitrogen stable isotope profiles for Ross seal males (**A**.) and females (**B**.)

**Table S1:**
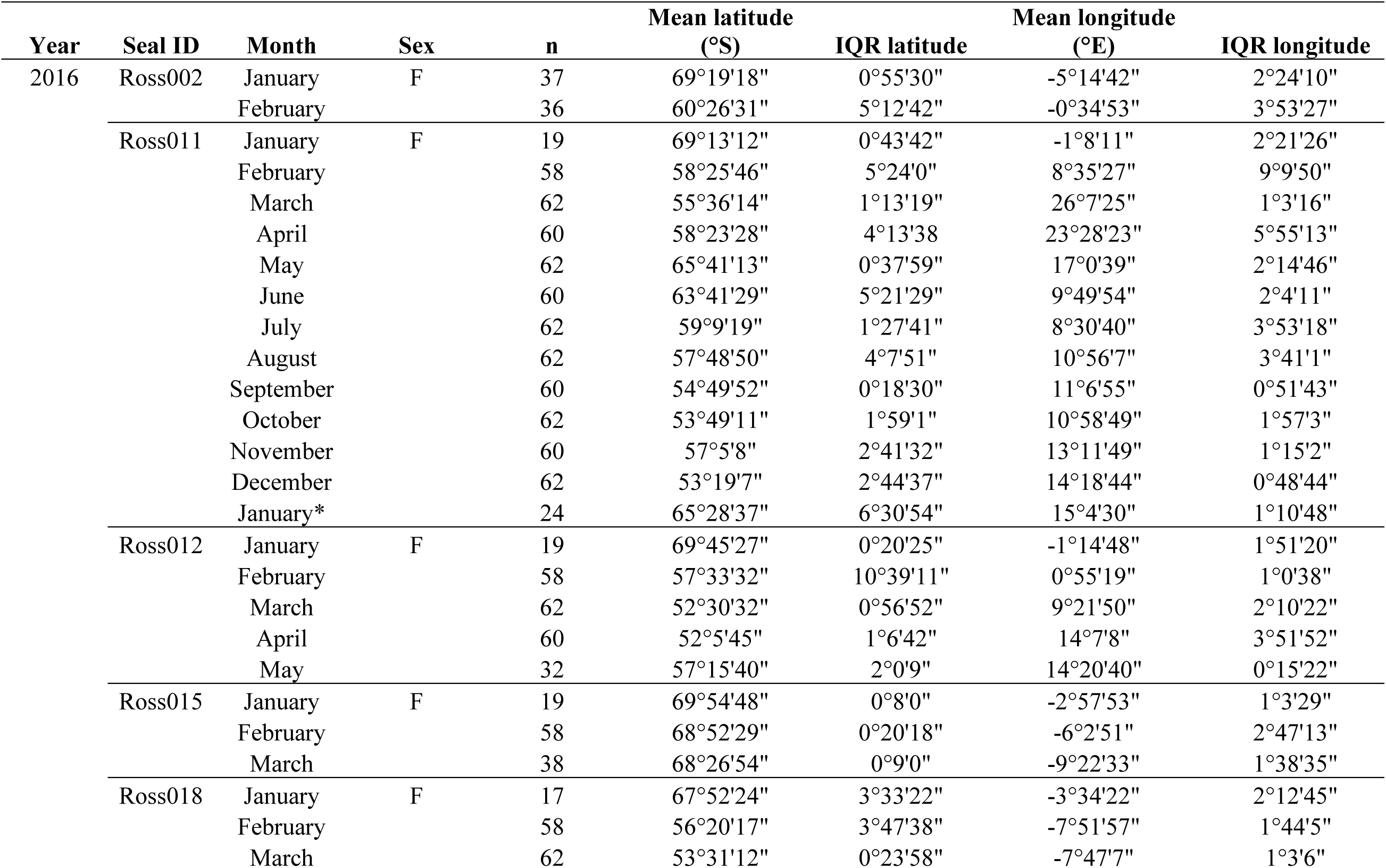
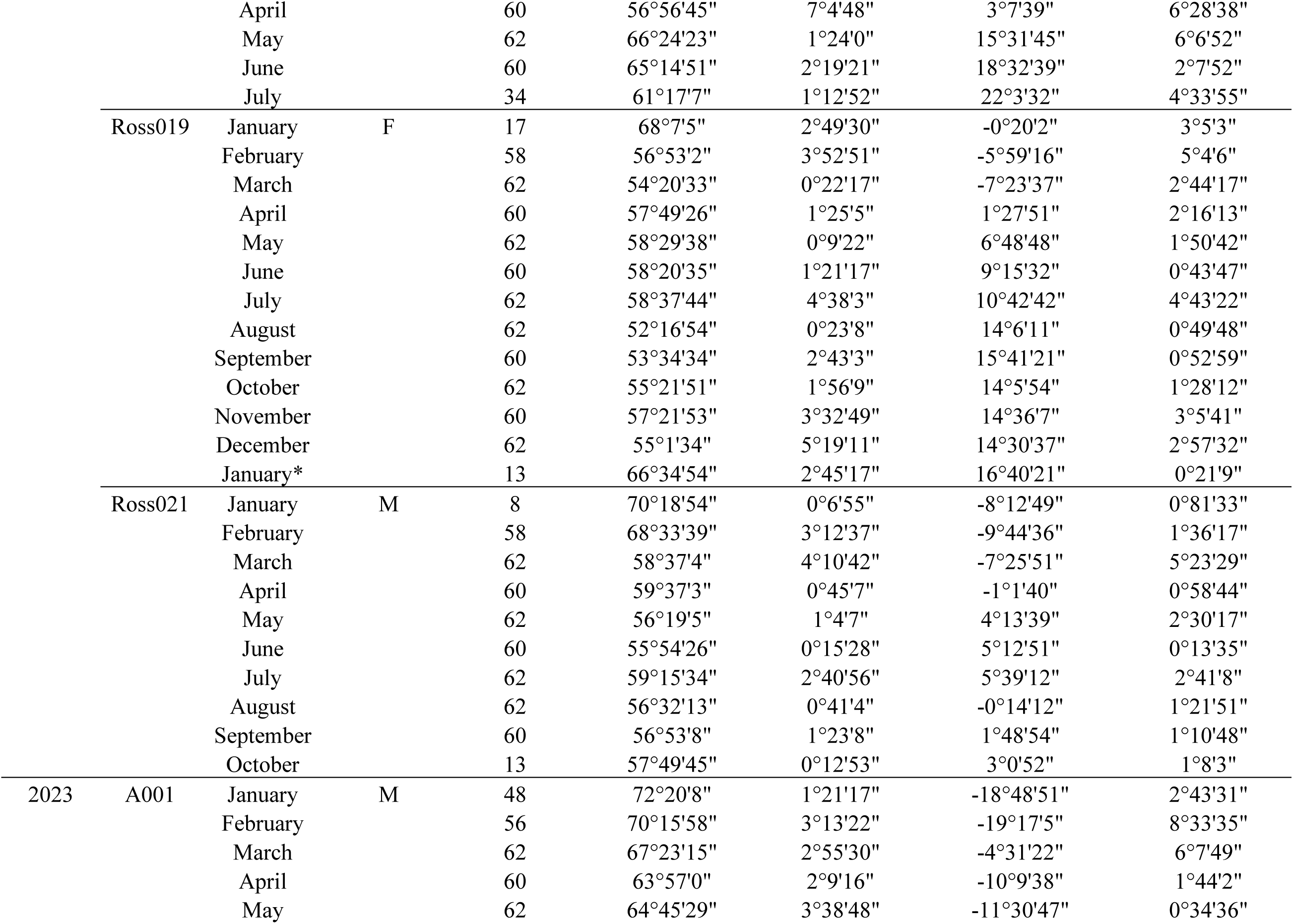
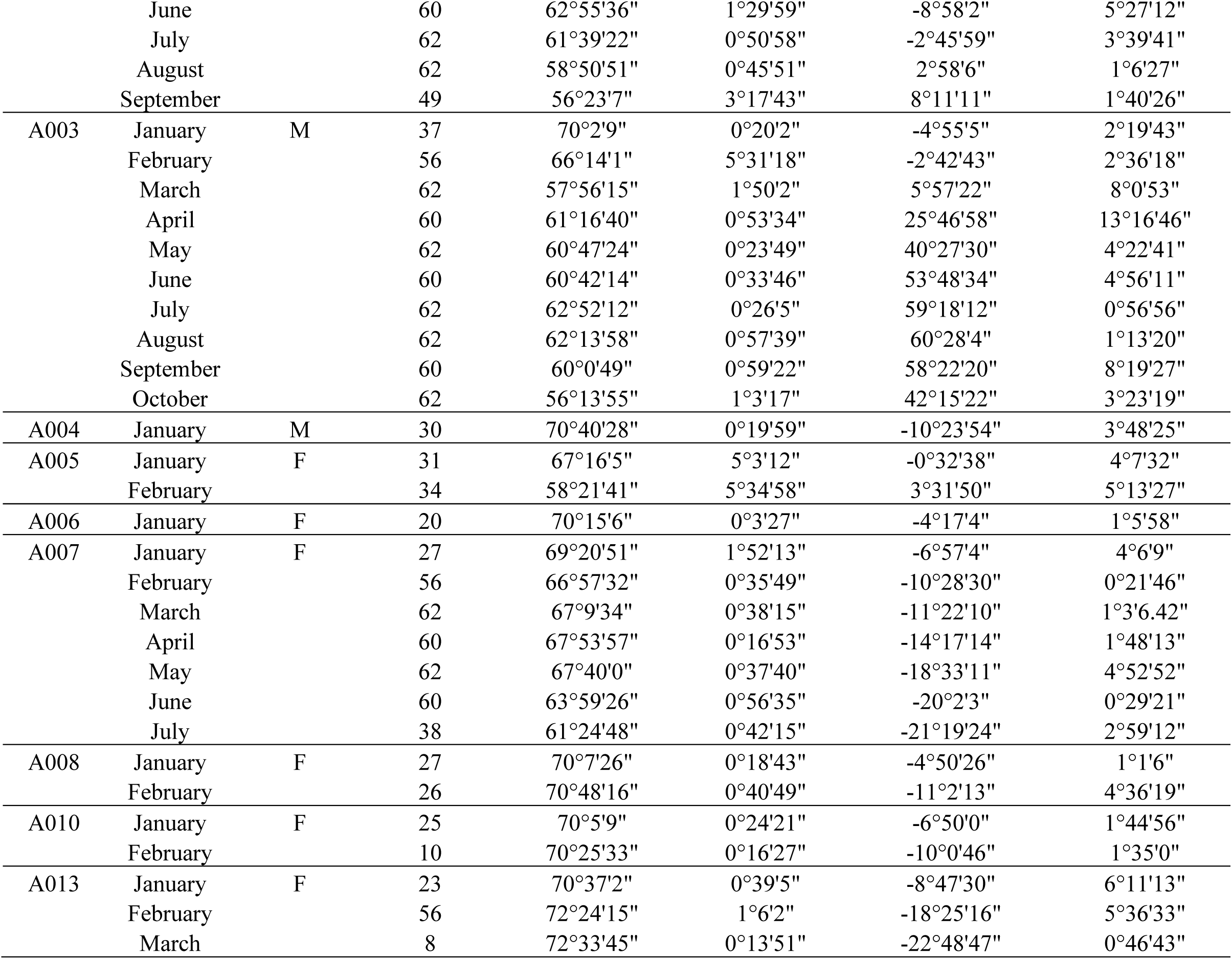
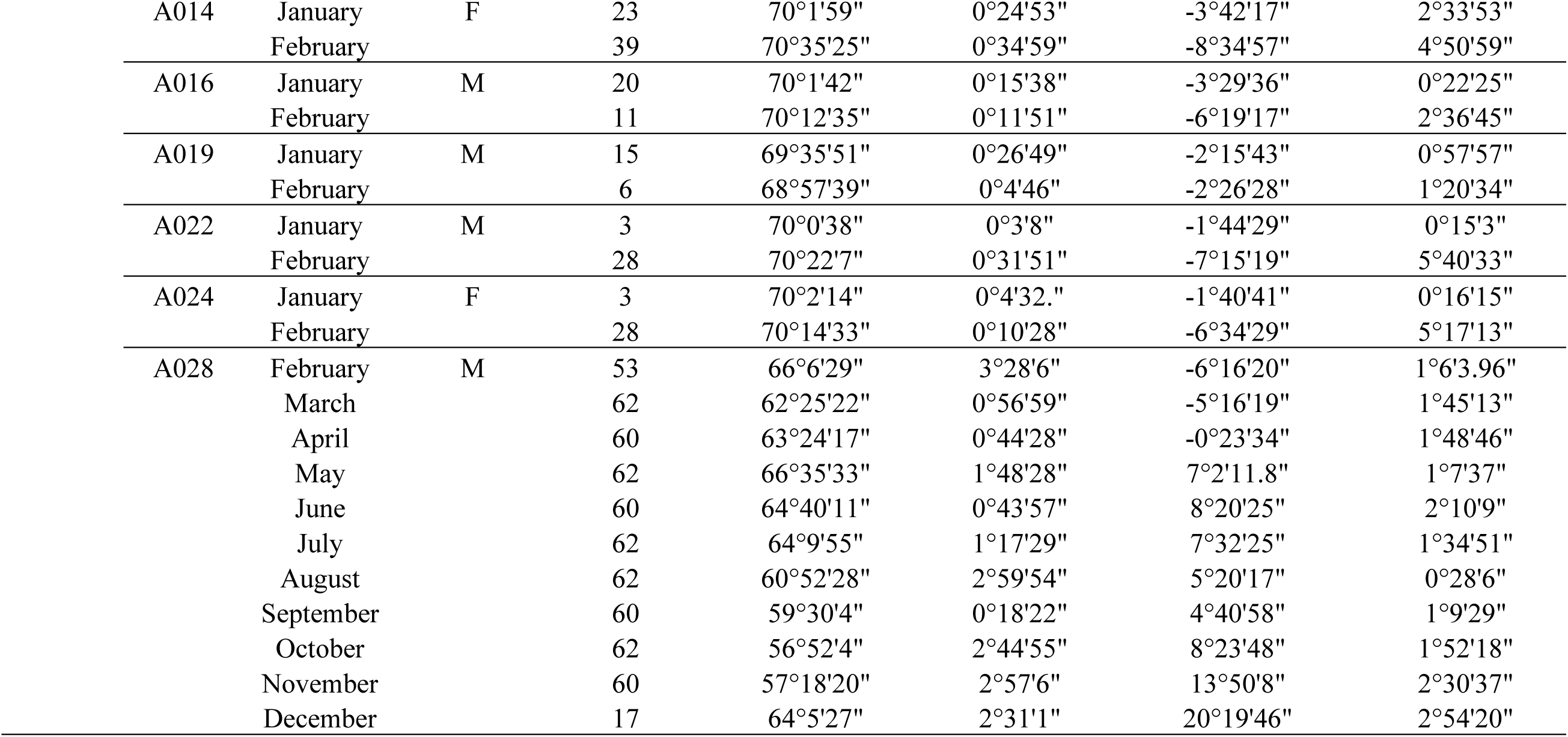
Latitudes and longitudes visited by seals. Abbreviations = n: number of tracking data after application of the 12 hour time step, IQR: Inter-Quantile Range

**Table S2:**
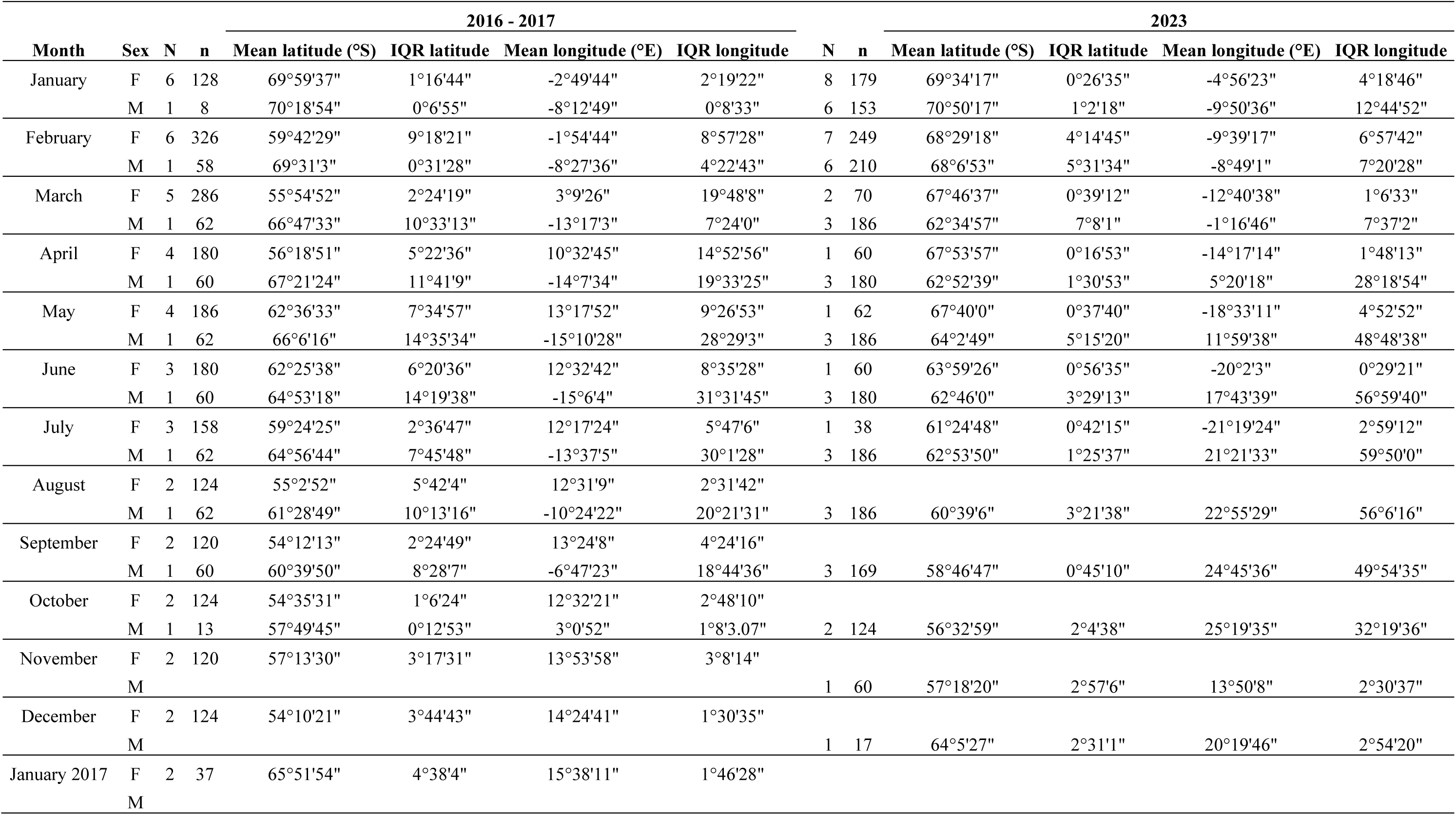
Latitudes and longitudes visited by seal sex. Abbreviations = N: number of seals, n: number of tracking data after application of the 12-hour time step, IQR: Inter-Quantile Range

**Table S3:**
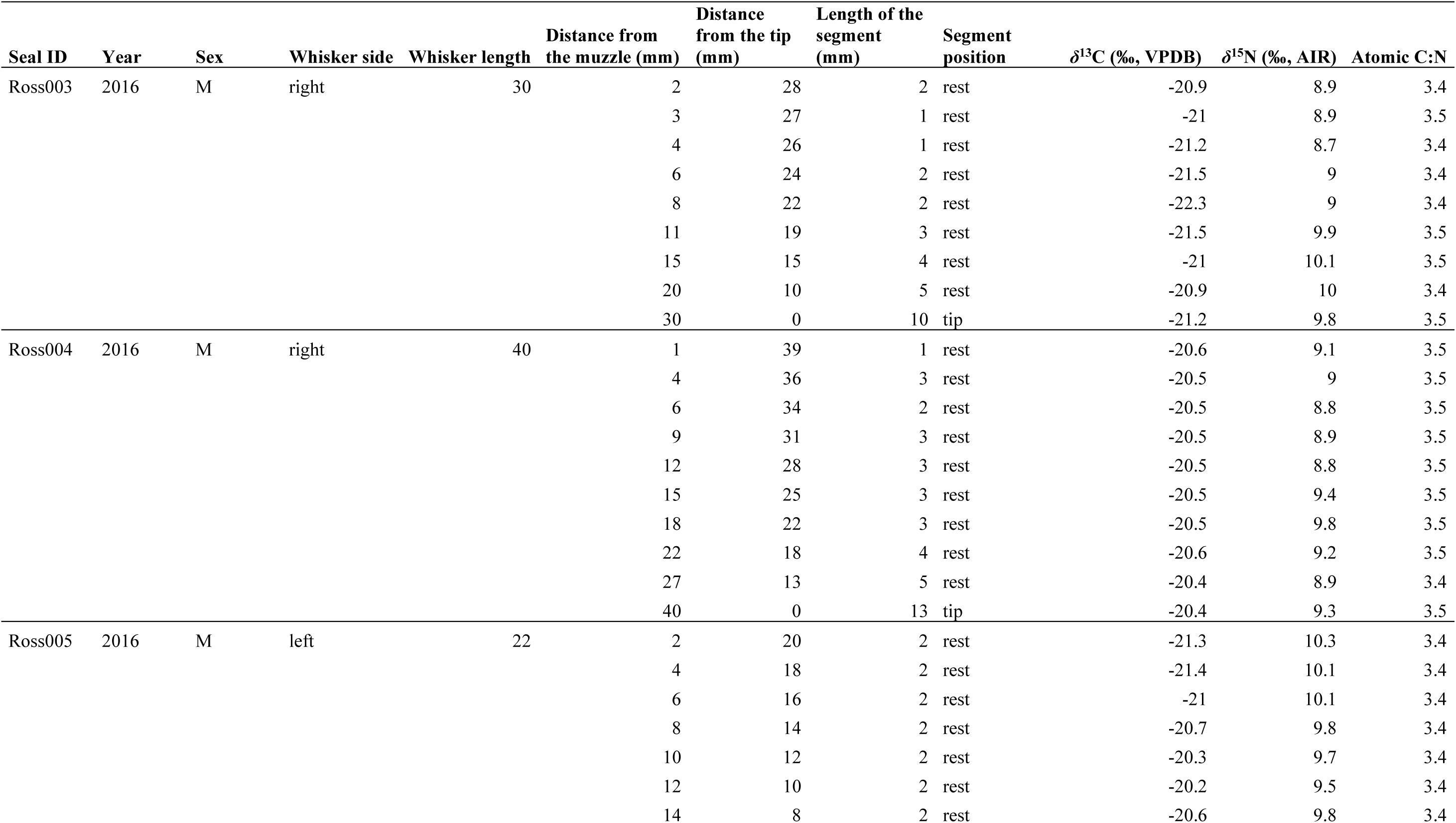
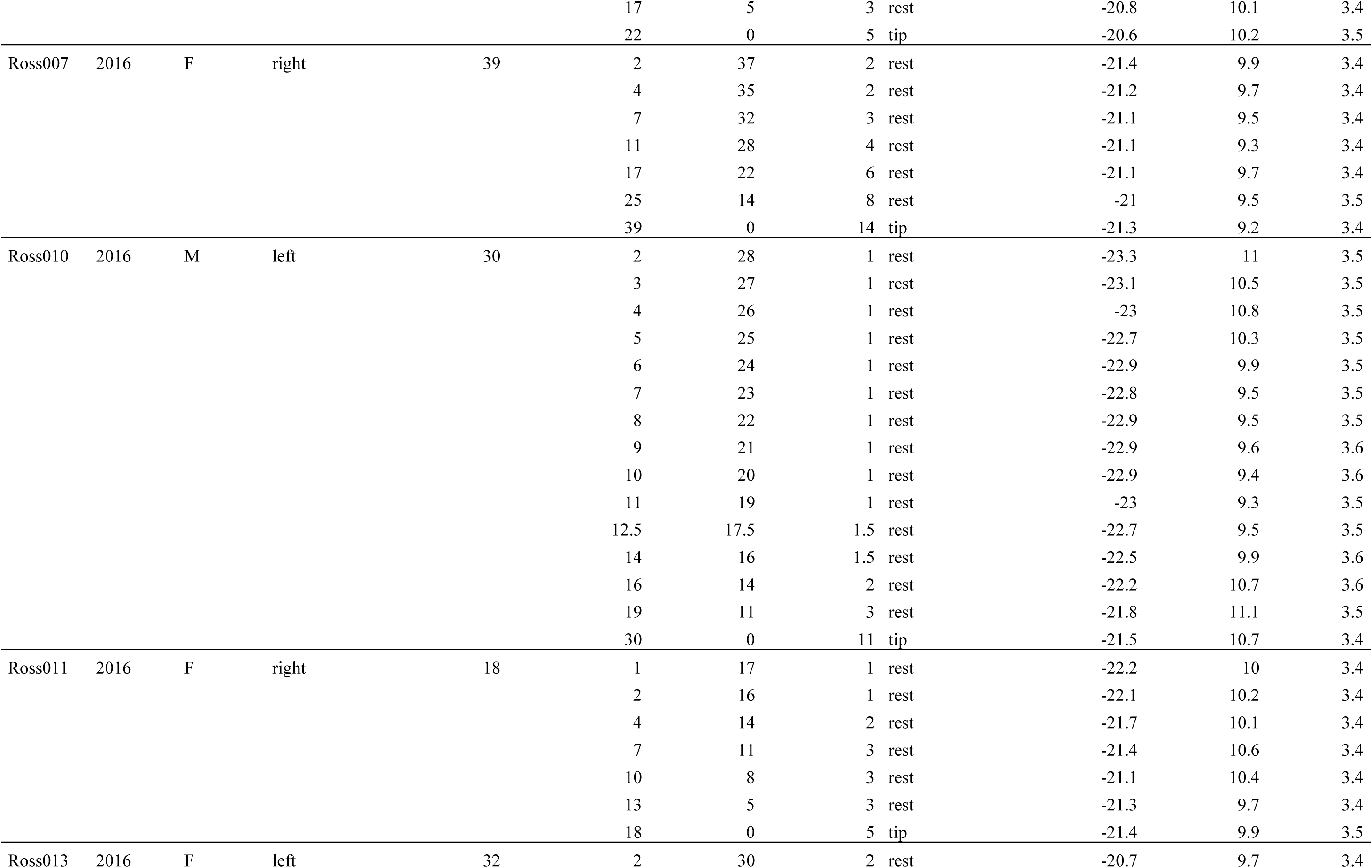
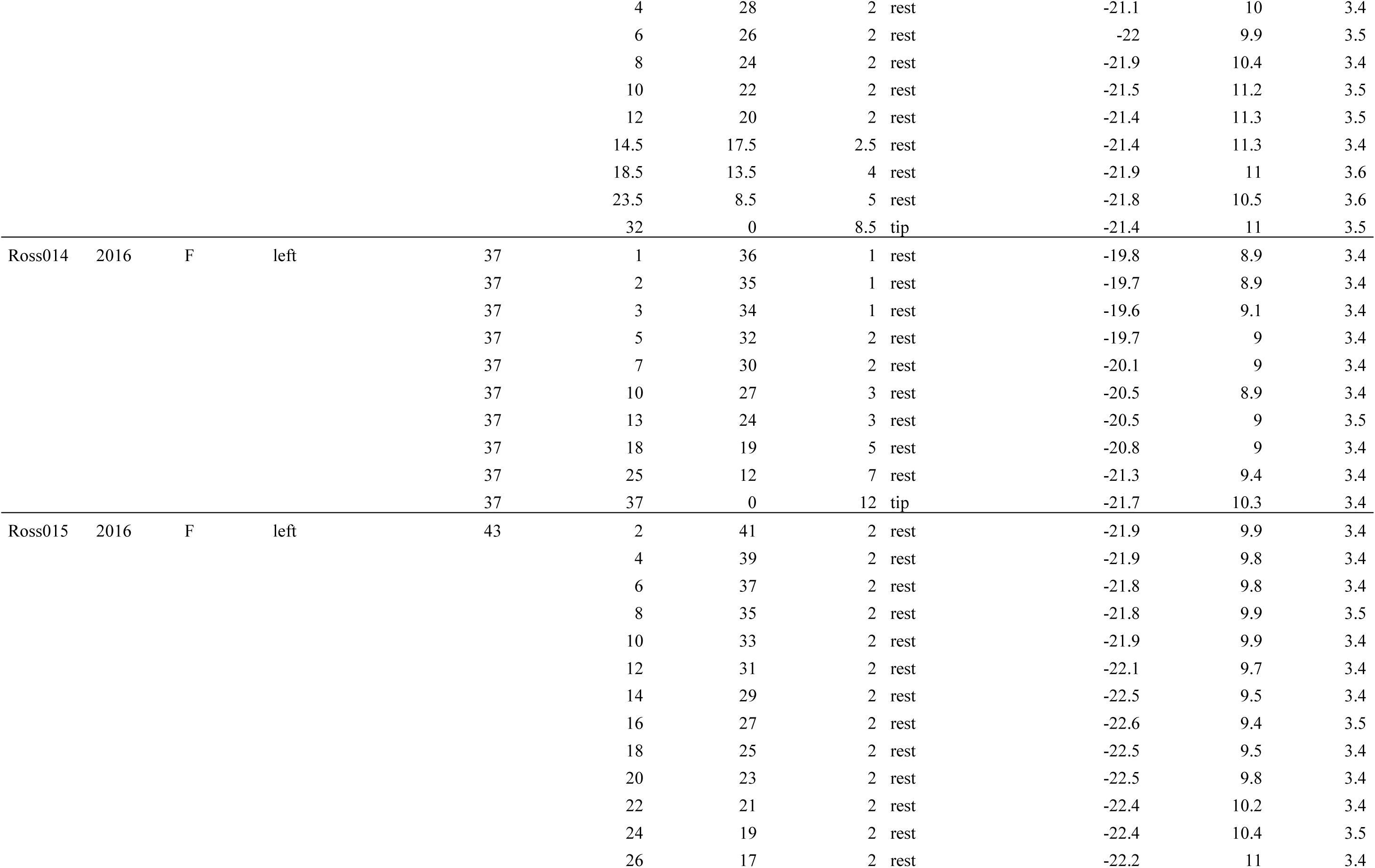
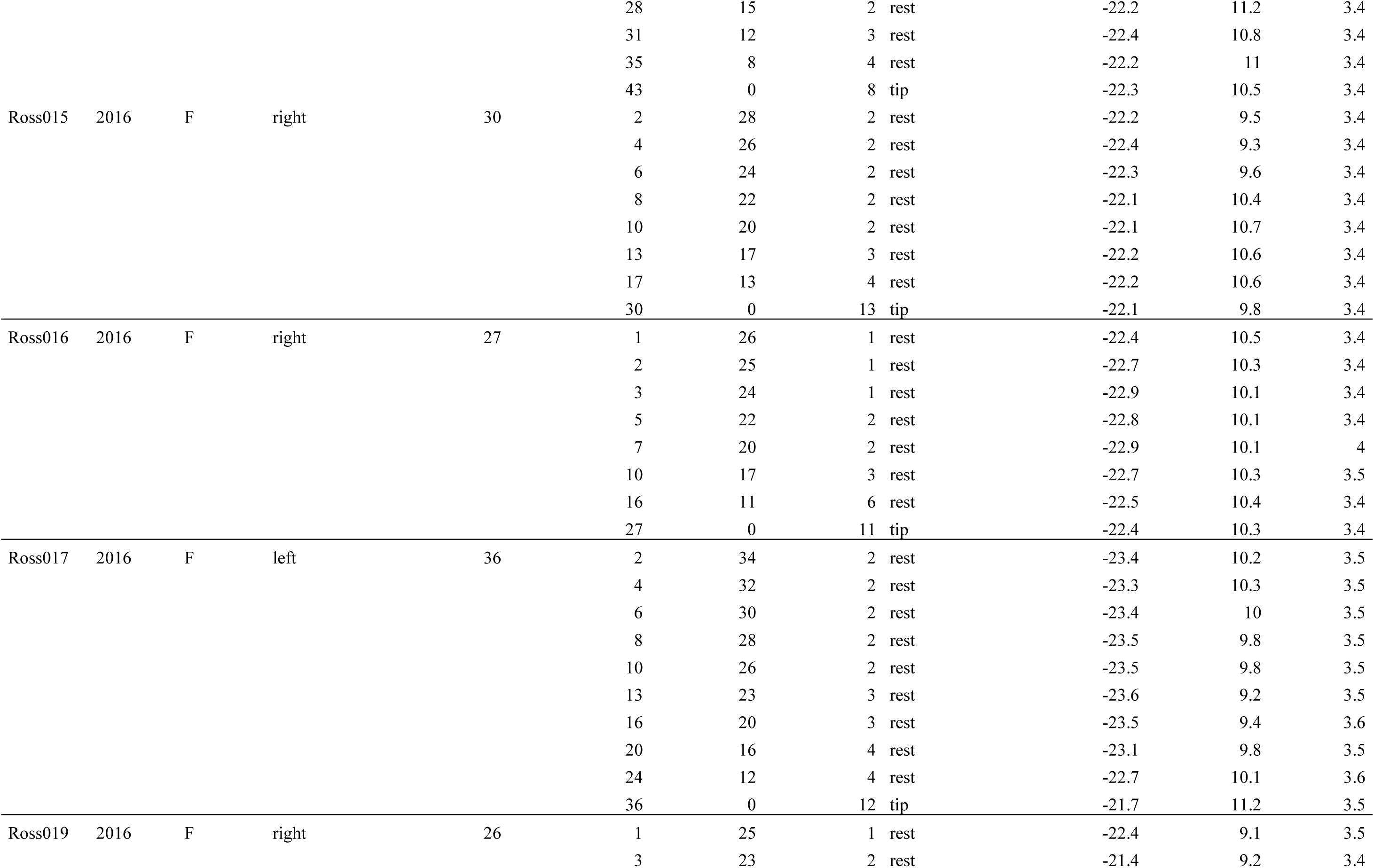
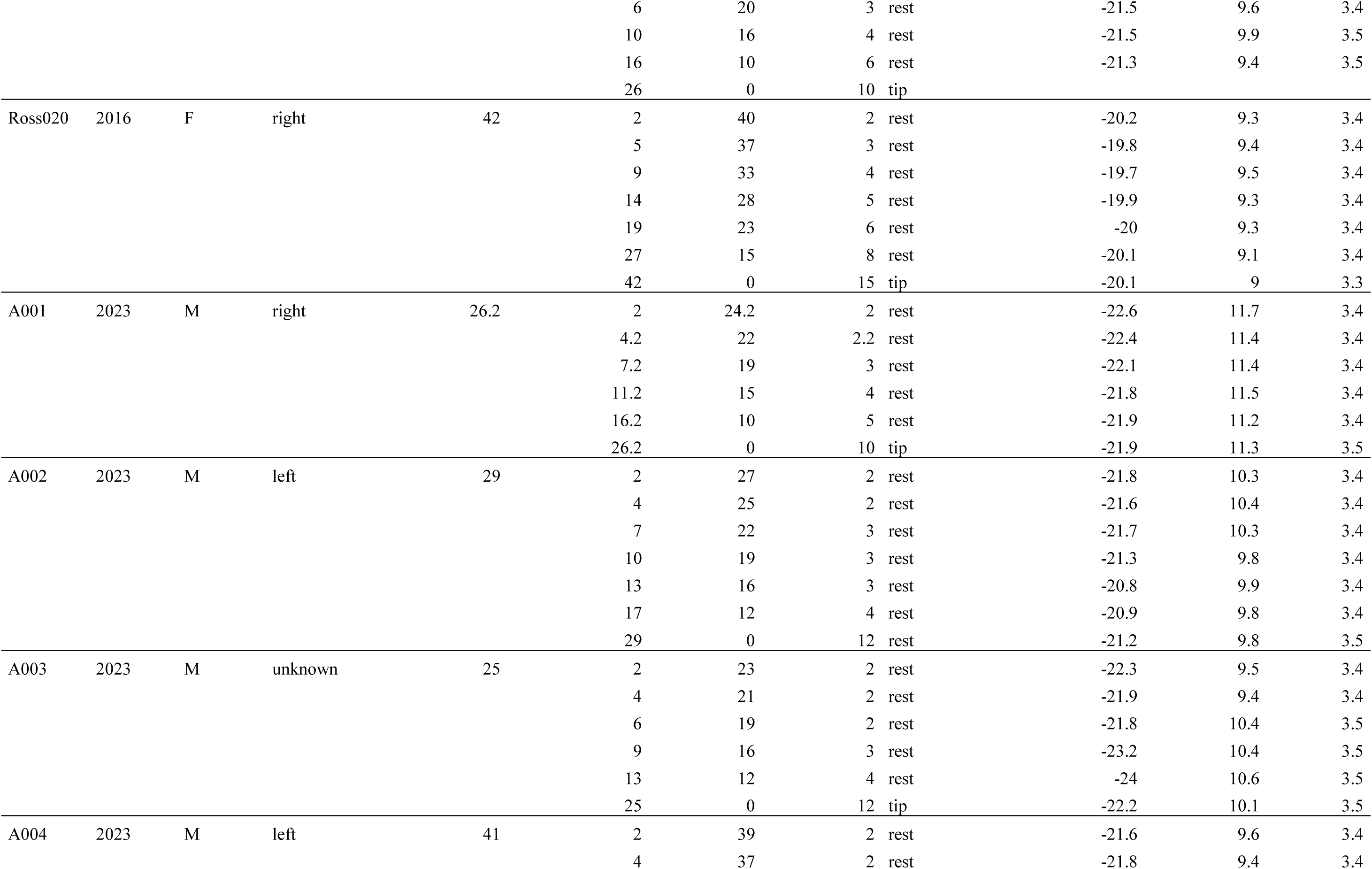
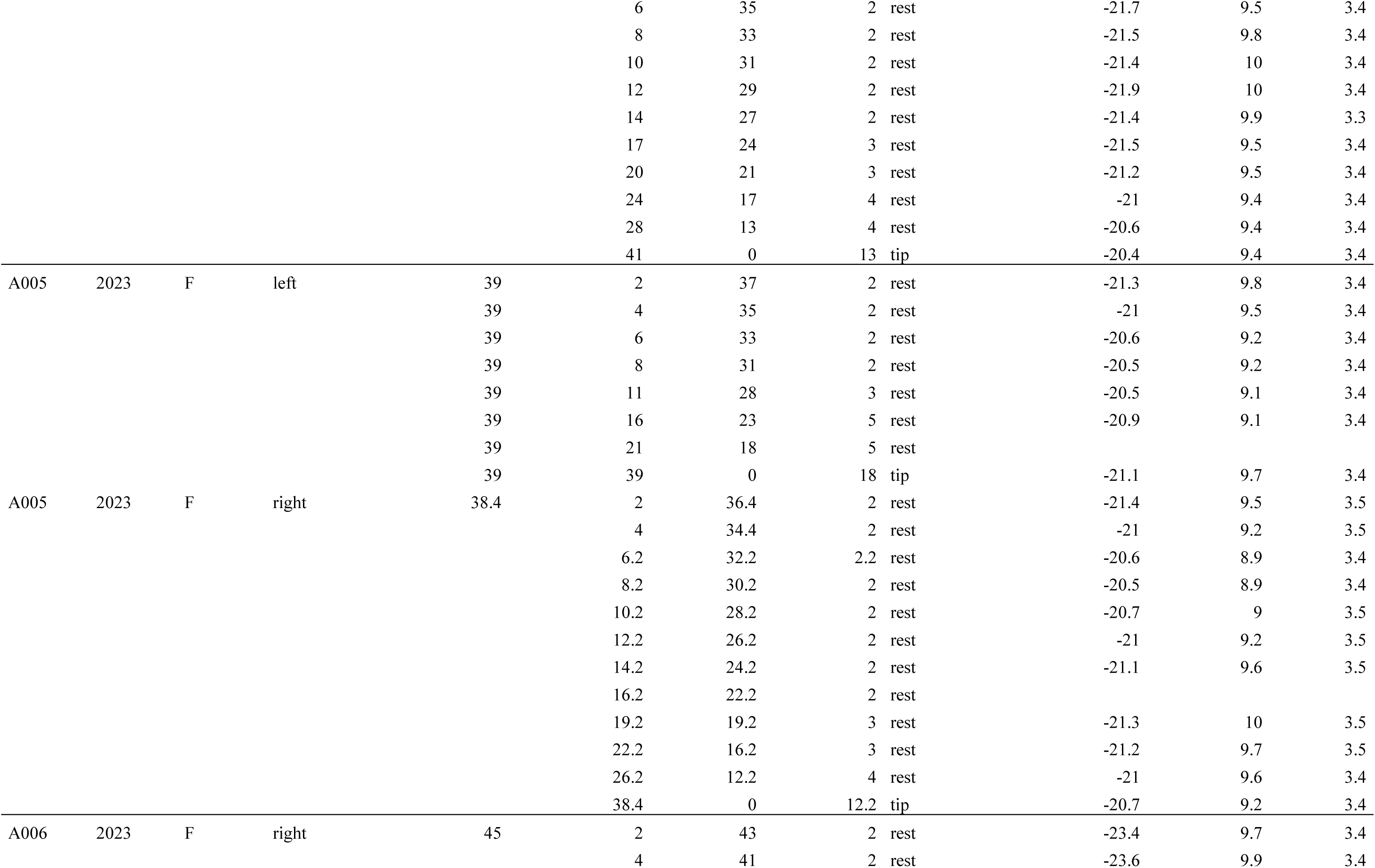
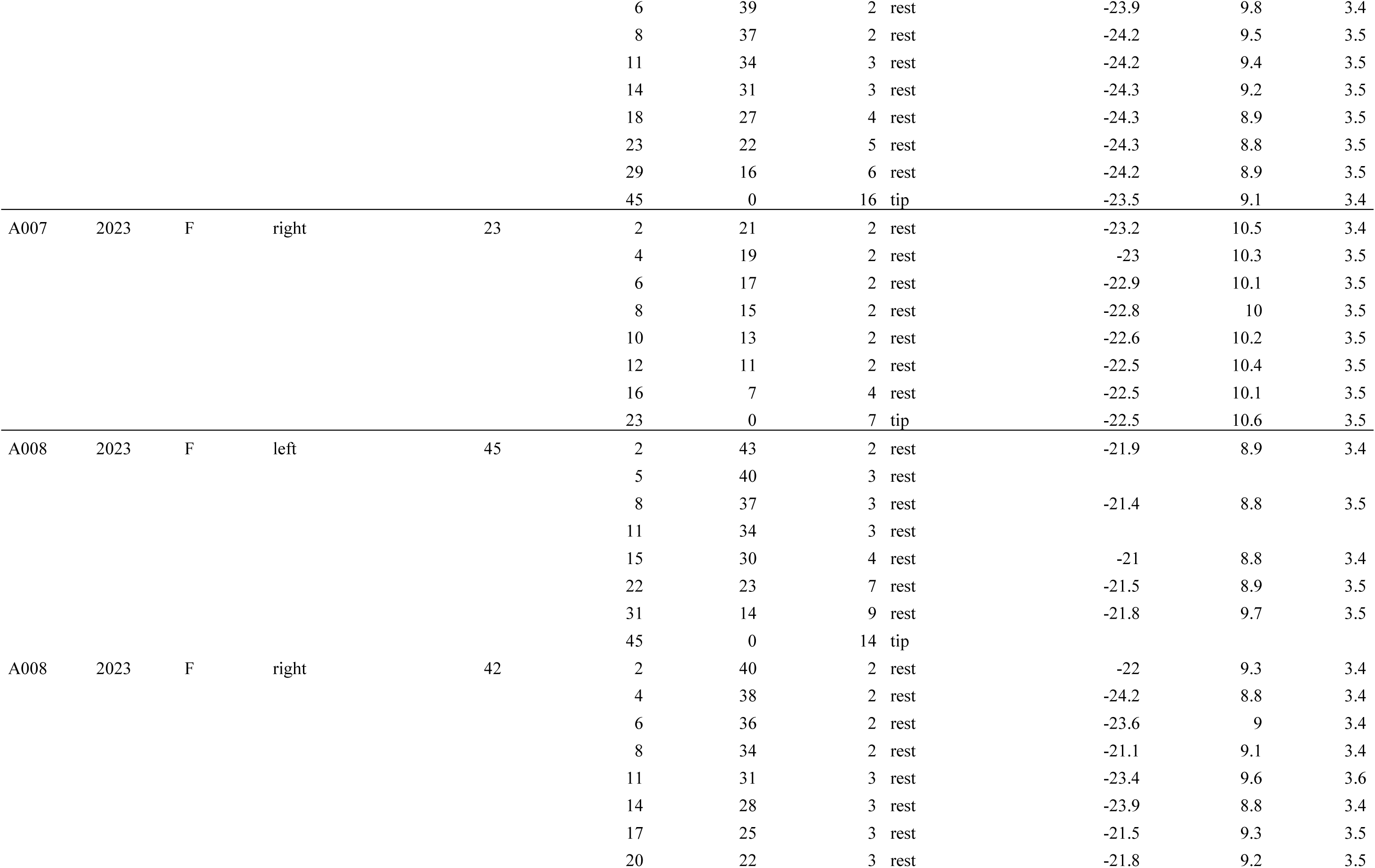
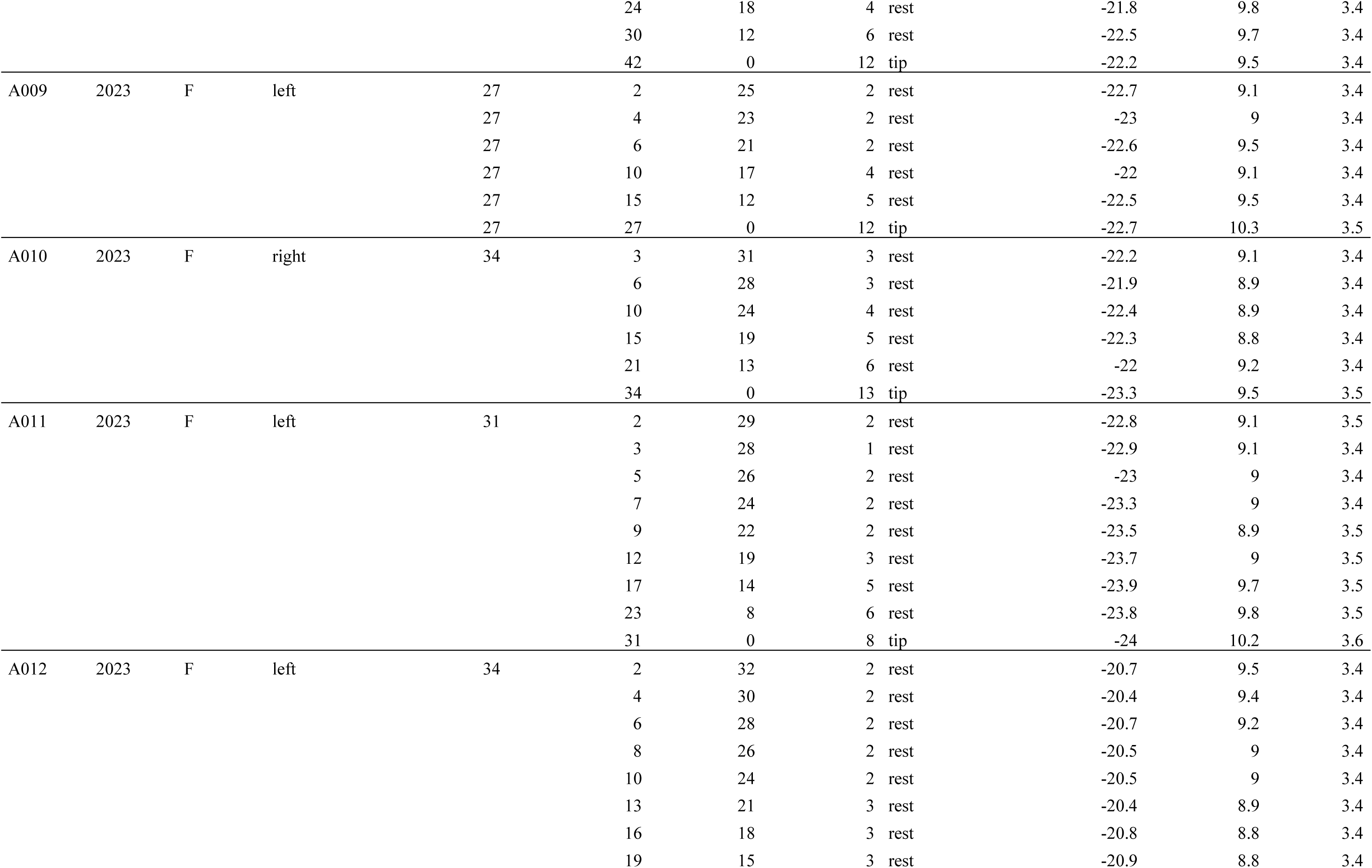
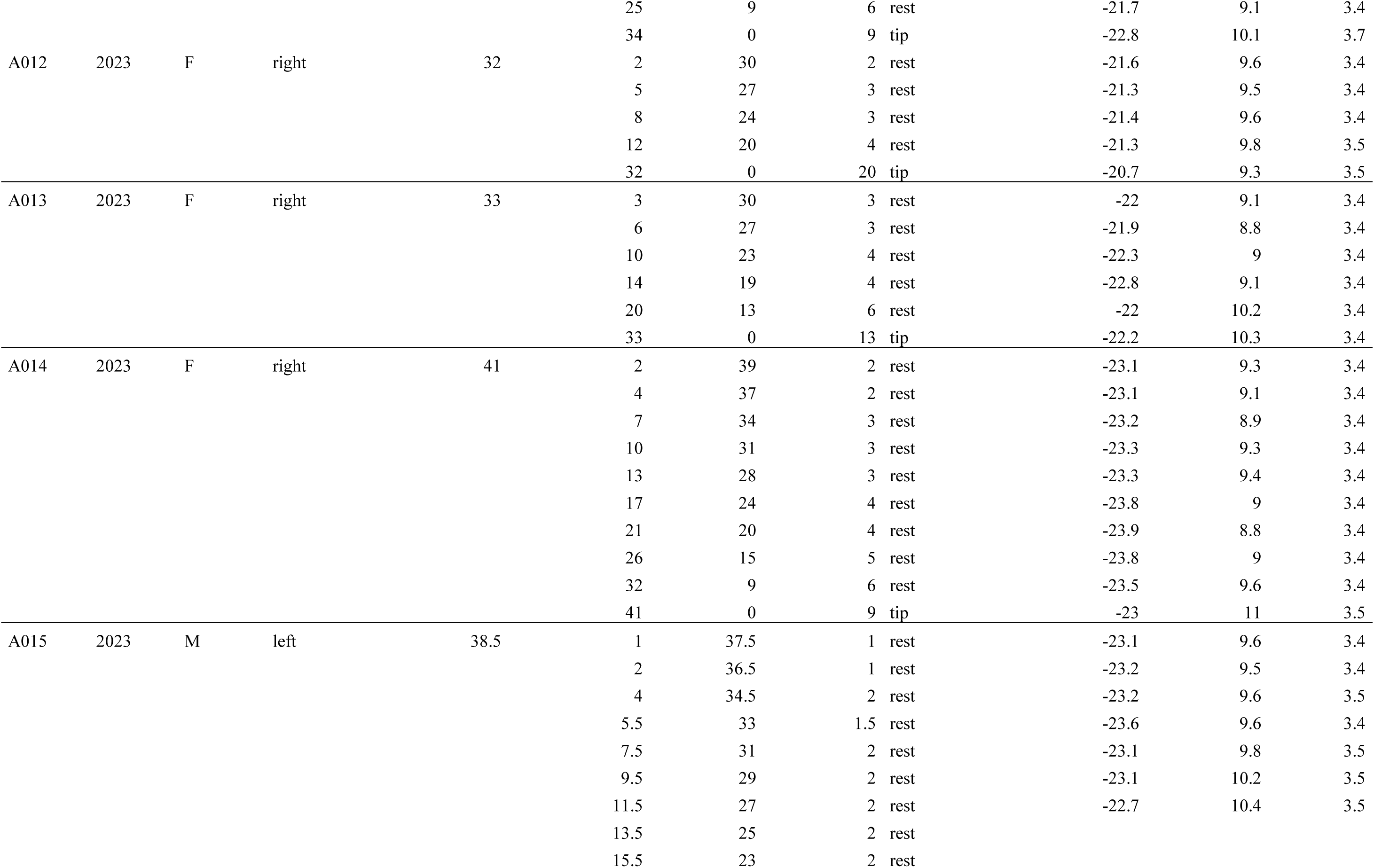
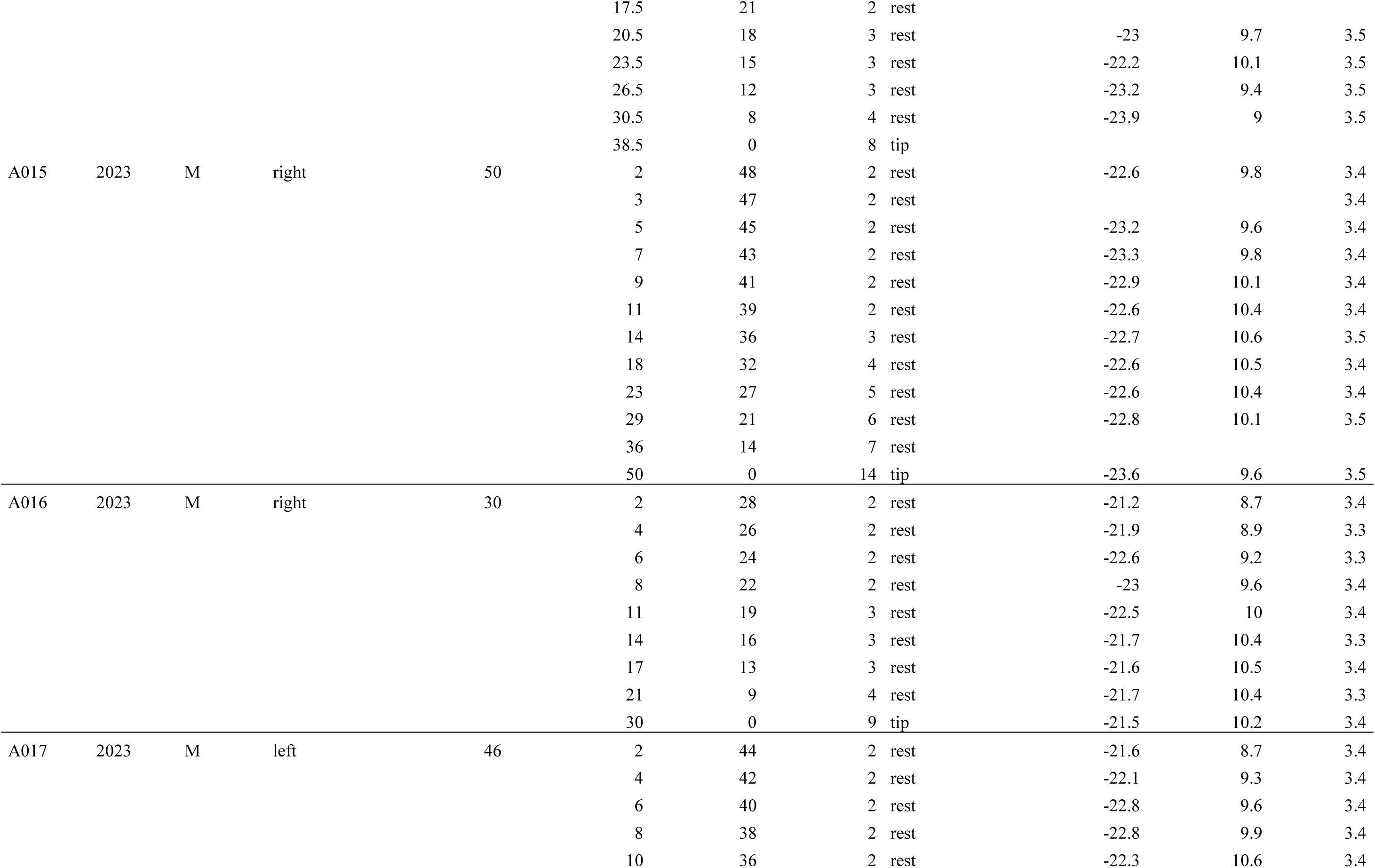
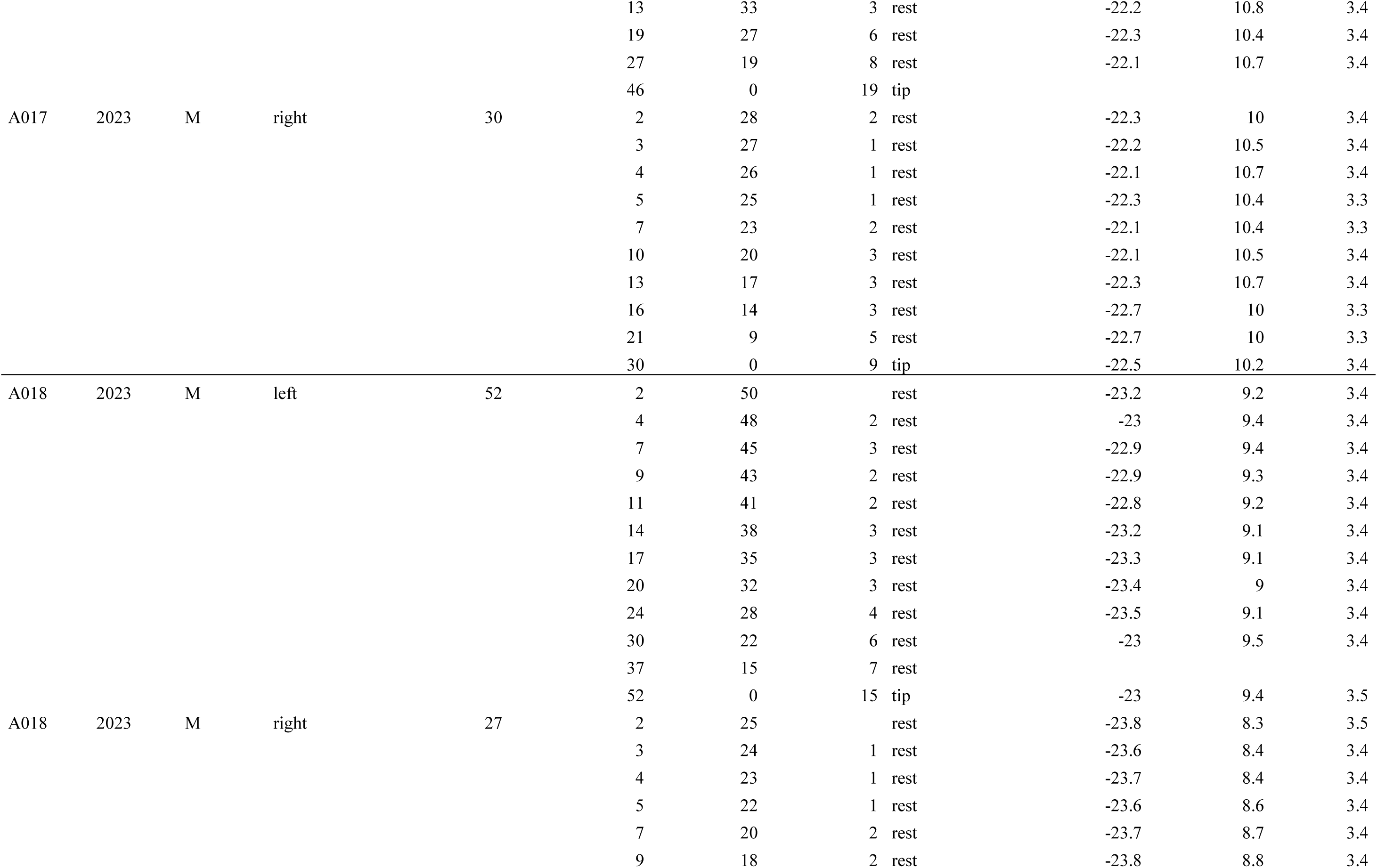
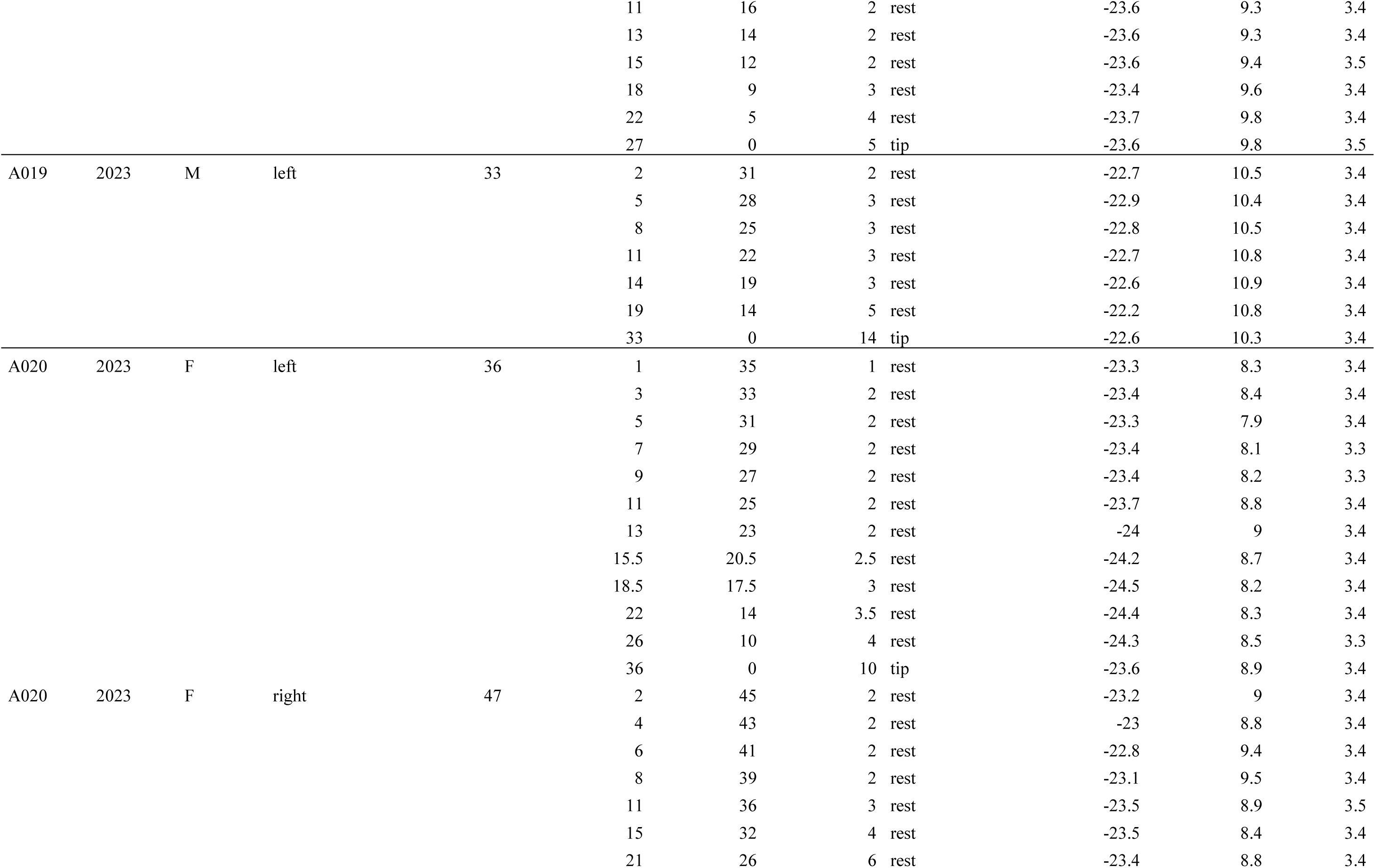
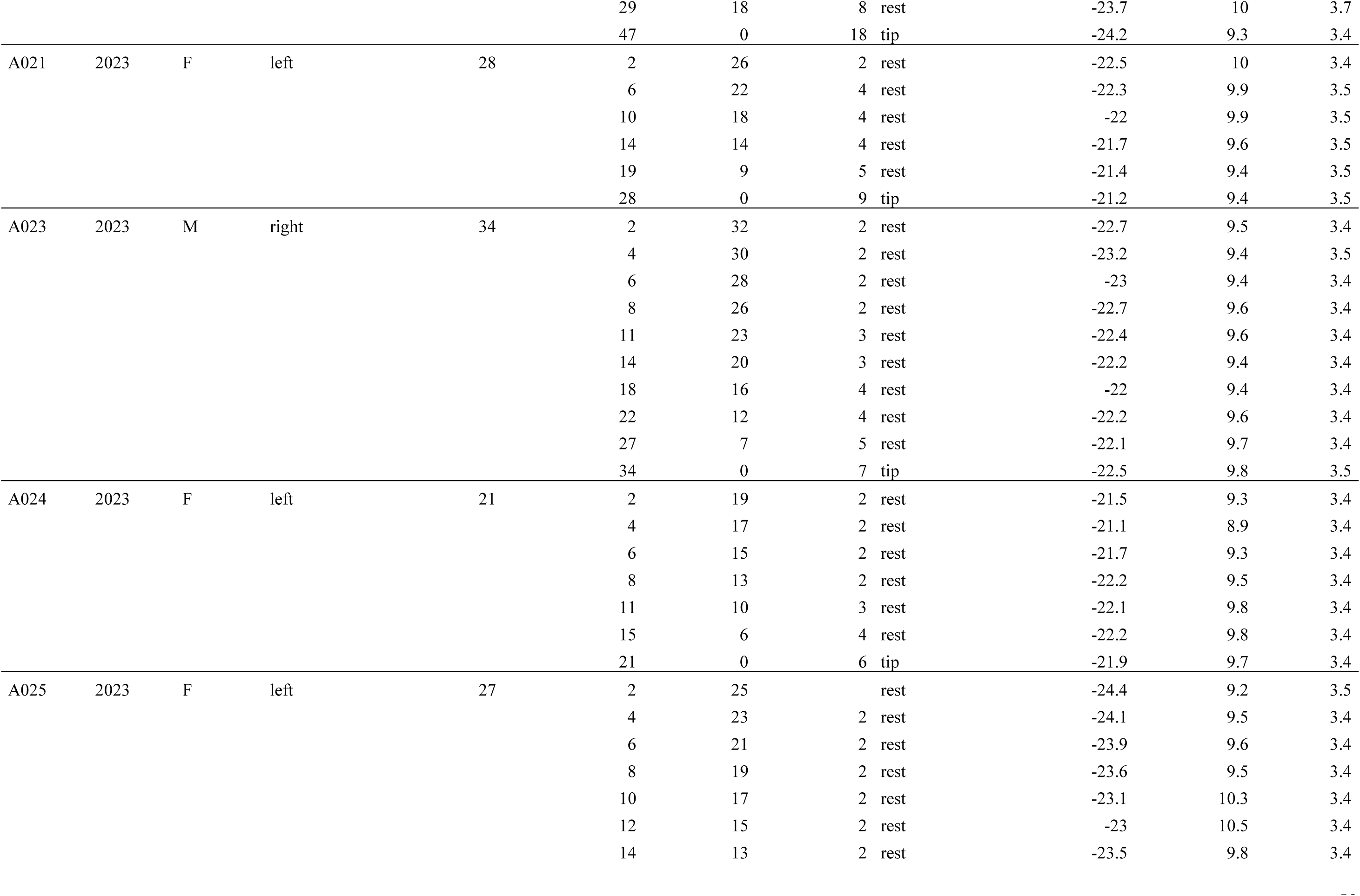
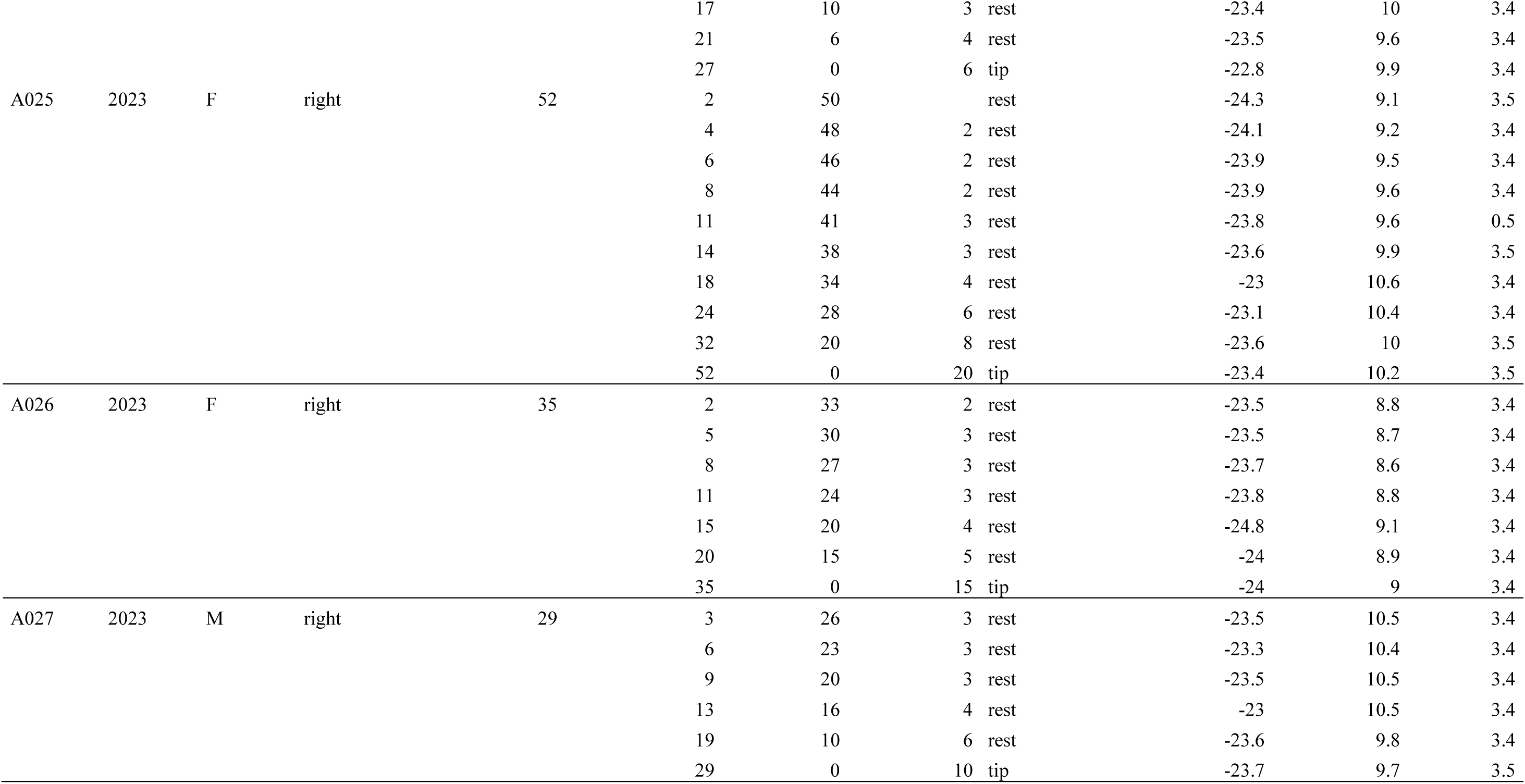
Detailed carbon (δ^13^C) and nitrogen (δ^15^N) stable isotope values for each whisker analysed.

